# Kaempferol over-accumulation in the flavonoid 3’ hydroxylase *tt7* mutant disrupts seed coat outer integument differentiation and compromises seed longevity

**DOI:** 10.1101/2022.03.08.483417

**Authors:** R Niñoles, P. Arjona, A. Hashim, E. Bueso, R. Serrano, I. Molina, J Gadea

**Author notes:** Author for correspondence: José Gadea.

## Abstract

- Seeds slowly accumulate damage during storage, which ultimately results in germination failure. The seed coat is the barrier between the embryo and the external environment, and its composition is critical for seed longevity. Flavonols accumulates in the outer integument, but the effect of altering flavonol composition on outer integument development has not been explored.
- Genetic, biochemical, ultrastructural and transcriptomics assays on a battery of loss-of-function mutants in the flavonoid biosynthesis pathway were used to study the effect of altered flavonoid composition in seed development and seed longevity.
- Controlled deterioration assays indicate that loss-of-function of the flavonoid 3’ hydroxylase TT7 gene dramatically affects seed longevity and seed development. Seed outer integument differentiation is compromised from nine days after pollination in *tt7* seeds, with a defective suberine layer and incomplete degradation of seed-coat starch. These distinctive phenotypes are not shared by other mutants showing also altered flavonoid composition. Double-mutant analysis indicate that over-accumulation of kaempferol is the primary cause of the observed phenotypes. Expression analysis suggest that the *tt7* flavonoid pattern affects transcriptional and non-transcriptional levels of regulation.
- The increase of kaempferols in the seed coat influences seed development. This positions TT7 as an essential player modulating seed coat development and seed longevity.

## INTRODUCTION

Gradual decrease of seed viability during storage is a major concern for agriculture. The seed coat is the physical barrier between the embryo and its external environment and improves viability by protecting the embryo from the entry of oxygen, which progressively damage seed constituents. The Arabidopsis seed coat originates from the ovule integuments, leading to a three-layered inner integument and a two-layered outer integument in the seed, each one following a distinct path during development (Haughn & Chaudhury, 2005). Early after fertilization, cells of the innermost layer of the inner integument (the endothelium) differentiates and synthesizes polymers of flavonoids called proanthocyanidins (PAs). These cells also produce an endosperm-surrounding lipidic cuticle (Loubéry et a 2018). At the desiccation stage, these cells die, and fused together with the two other layers of the inner integument, constituting the brown pigment layer (BPL). The two layers of vacuolated cells of the outer integument are initially indistinguishable. At 7 DAP, cells of both layers accumulate starch granules, that will degrade at later stages to reinforce cell walls. Then, they diverge in fate. The external epidermal layer synthesizes mucilage (a pectinaceous carbohydrate) and secretes it to the apoplast, forcing the cytoplasm to the center of the cell, and leaving at maturity a granule-free cytoplasmic volcano-shaped column (the columella). In the inner layer of the outer integument, flavonols (but not PAs) are synthesised (Xuan et al, 2018). Later on, these cells synthesize suberine, and finally die and crush together, constituting the suberized palisade layer (Haughn & Chaudhury, 2005).

Different compartments of the seed coat are known to contribute to longevity. Specialized in sealing off specific plant tissues, lipid polyester barriers are determinants for seed viability. Mutants in genes required for lipid polyester biosynthesis renders highly permeable seed coats, (Beisson et al., 2007; De Giorgi et al, 2015), correlating with decreased longevity Renard et al., 2020), and accumulation of these biopolymers have the opposite effect (Renard et al, 2021). The influence of flavonoids to seed longevity is more controversial. Initially, Debeaujon et al, 2000 observed sensitivity to ageing in a set of mutants deficient in the flavonoid biosynthesis pathway (*transparent testa, tt*), and proposed for these compounds a role as seed antioxidants. Recently, Lobéry et al, 2018 demonstrated that PAs act as structural components of the endosperm-associated cuticle, and in mutants defective in PAs biosynthesis this cuticle is affected or absent. This would explain their high seed permeability, which would compromise seed longevity. Seed coat flavonols have received less attention, and whether they influence development of the outer integument is not known.

Flavonols are well-known inhibitors of protein-kinase activity, some of them essential for plant and seed development. Flavonols bind to phosphoinositide-kinases (Lee et al, 2010; Kong et al, 2011), small GTPases (Sharma et al, 2019) glucosidases (Peng et al, 2016) or actin (Böhl et al, 2008), thus interfering or modulating cellular metabolism and communication. Inhibition of auxin transport mediated by phosphoinositide-kinases is one of the most studies effects of flavonols (Peer and Murphy, 2007; Ischebeck et al, 2013), but given their multiple consequences on enzymatic activities, additional ones are also possible. Cell-cell communication through plasmodesmata (PD) coordinates distribution of sugars and the development of many processes by means of mobile signals that are selectively transported to target cells to specify their function (Han et al, 2014; Han & Kim, 2016; Liu et al, 2018). In the seed coat, the outer integument forms a symplastic domain connected to the funiculus, which provides assimilates and signals for proper differentiation (Stadler et al, 2005). Plasmodesmal permeability regulation seems then essential for seed coat development. Whether flavonols of the outer integument affect this process is unknown.

In this study, we show that the loss-of-function of the flavonoid 3’ hydroxylase *TT7* (*TRANSPARENT TESTA 7*) gene dramatically affects seed longevity. In *tt7* mutants, seed coat differentiation is compromised, leading to dry seeds with severe outer integument defects, such as a defective suberized palisade layer and incomplete seed-coat starch degradation. Mutants defective in seed-coat starch degradation present reduced germination after ageing treatments, revealing a novel aspect contributing to this trait. These *tt7* defects are seed-coat specific and are not shared by other flavonoid biosynthetic mutants. Kaempferol overaccumulation in *tt7* seed coat is causing these phenotypes. Transcriptomic analysis suggests altered expression in *tt7* seeds of suberine and starch metabolism genes, as well as genes involved in cell signalling, probably as consequence of kaempferols inhibiting key enzymes in these pathways. These findings show that changes in flavonol composition can have a tremendous impact on cellular processes involved in seed coat development. Given that many abiotic stresses alter flavonol composition, understanding the impact of these changes is of paramount importance to obtain high quality seeds.

## MATERIALS ANS METHODS

### Plant material and growth conditions

*tt7-7* (Col-0 background) was obtained from the Nottingham Arabidopsis Stock Centre (NASC); *tt3-4, tt4-8, tt7-4, tt10-2, ban-1* and *fls1-3* in Wassilevskija (WS) background and *tt5-1* in Lansdberg *erecta* (Ler) background were kindly provided by Dr. Isabelle Debeaujon. For auxin localization experiments, the DR5:GFP marker line was used (Weijers *et al* 2006).

*Arabidopsis thaliana* seeds were surface sterilized with 70% ethanol 0.1% Triton X-100 for 15 m and rinsed three times with sterile water. Seeds were stratified for 3 days at 4 °C. Germination was carried out on plates containing Murashige and Skoog (MS) salts with 1% (w/v) sucrose, 10 mM 2-(N-morpholino) ethanesulfonic acid and 0,9% (w/v) agar, pH was adjusted to 5.7. To obtain seeds, plants were grown under greenhouse conditions (16 hr light/8 hr dark, at 23±2 °C and 70 ± 5% relative humidity) in pots containing a 1:2 vermiculite:soil mixture. Control and mutant plants were grown simultaneously and seeds were collected and stored under the same conditions.

When indicated, a naphthylacetic acid (NAA) treatment was performed by spray application of a 50µM NAA + 0,02% Silvett solution to Col-0 developing siliques (6-10 days after pollination, DAP). 30 m after the treatment, siliques were harvested.

### Artificial ageing assays

For Controlled Deterioration Treatment (CDT), seeds were maintained for 14 days (17 days for reciprocal crosses) at 37 °C at 75% RH (Righetti et al, 2015). The Elevated Partial Pressure of Oxygen assay (EPPO) (Groot et al., 2012) was performed for 5 months at 5 bar O2 with a 40% RH. Ageing treatments were performed prior to seed sterilization and stratification.

### Tetrazolium permeability and reduction assays

Triphenyltetrazolium salt penetration assay was performed as described by Molina et al, 2008. Briefly, seeds were incubated in the dark in 1% (w/v) tetrazolium red at 30 °C for 8 to 72h as indicated. Then, formazan was extracted and quantified as absorbance at 485 nm. For tetrazolium salt reduction assay, seed coats were gently broken and seed coats and embryos were soaked in 1% tetrazolium solution for 1day in the dark at 30°C. Images were taken with a Leica DMS1000 macroscope.

### Biochemical assays and transmission electron microscopy experiments

Mucilage staining of dry seeds was performed as described in Western et al, 2001. Lipid polyester staining was performed as described in Beisson et al. (2007). For starch staining, seeds or rosettes (18 days after sowing) were boiled in 70% EtOH for 5 min. After washing with water, iodine solution was added and after 5 m images were taken with a camera (rosettes) or Nikon Eclipse E600 microscope (seeds). Transmission electron microscopy were performed as described in Supplemental Methods.

### Seed polyesters analysis

Seed samples were delipidated and dry residues were depolymerized by methanolysis in the presence of sodium methoxide. After acetylation of the CH_2_Cl_2_-extractable products, monomers were analysed by gas chromatography according to Supplemental Methods.

### CRISPR/Cas9 vector construction for *tt7* editing

Two sgRNAs were designed to generate *tt7* loss-of-function mutations (Supplemental Methods). These sgRNAs were PCR-amplified with the pCBC-DT1T2 vector using the primer sets listed in Supplementary Table 3. PCR products were digested with BsaI and subcloned into the pHEE401 binary vector containing *Zea mays* codon-optimized Cas9 under the control of an egg cell-specific promoter (Wang et al. 2015). Positive constructs were introduced into *Agrobacterium tumefaciens* GV3101 and used for transformation of *tt3-4* or *fls1-3* mutants by floral-dip method. Identification of *tt7* edited plants was performed according to Supplemental Methods.

### DR5-driven fluorescence analysis

Developing siliques of DR5:GFP or DR5:GFP x *tt7* lines where fixed with 4% paraformaldehyde in 1 x PBS for 1 h at 22°C. Fixed tissues were washed twice with 1 x PBS, transferred to ClearSee solution and cleared at room temperature with gentle agitation for 7 days. Then, seeds where separated from siliques and mounted on slides with ClearSee solution for imaging. A confocal ZEISS-LSM710 microscope was used. Laser used was Argon with an excitation lambda of 488 nm, and an emission spectrum of 500-550 nm was registered.

### RNA extraction and analysis

Seed total RNA extraction was performed according to Oñate-Sánchez and Vicente-Carbajosa (2008). DNA removal was performed with DNAse kit (E.Z.N.A). About 1 μg RNA was reverse-transcribed using the Maxima first-strand cDNA synthesis kit (Thermo Fisher Scientific). qRT-PCR was performed with the PyroTaq EvaGreen qPCR Mix Plus (ROX; Cultek) in a total volume of 20 μl using an Applied Biosystems 7500 Real-Time PCR System. Data are the mean of three replicates. Relative mRNA abundance compared to PP2AA3 or AT5G55840 (Czechowski *et al*,2005) was calculated, using the comparative ΔCt method. Primers for qRT-PCR are listed in Supplemental table 3. For RNA seq, RNA was extracted from developing seeds (8 DAP). Twenty-million paired-ends 50pb reads per library were sequenced. Three replicates were used per genotype. Data analysis was performed according to Supplemental Methods.

## RESULTS

### The seed coat-based reduced longevity of *tt7* seeds is distinctive among flavonoid mutants

Debeaujon et al, 2000 reported a reduced resistance to seed ageing in a set of flavonoid mutants, but the contribution to this trait of the different subgroups of flavonoids was not explored further. We assayed seed longevity in a set of selected mutants whose metabolomic profiles are available. *tt4-8* and *tt5-1* do not contain any kind of flavonoids in their seeds, whereas *tt10-2* seeds accumulate more quercetin-derived flavonols and more PAs (cyanidin-derived, the type present in the wild-type Arabidopsis seed). *ban-1* and *tt3-4* seeds do not contain PAs, whereas flavonols are absent in FLAVONOL SYNTHASE (FLS) mutant seeds (*fls1-3*). Finally, FLAVONOID 3’-HYDROXYLASE mutant (*tt7-4*) seeds do not contain quercetin-derived flavonols, but accumulate large amounts of kaempferol-derived flavonols and contain an unusual composition of PAs (epiafzelechin-derived) (Pourcel et al, 2005, Routaboul et al, 2006; Schulz et al, 2016).

Artificial ageing assays reveal a very different response among these mutants. To magnify differences, ageing conditions were adjusted so that wild-type seeds were able to germinate up to 60% in controlled-deterioration treatment (CDT) and 90% in Elevated Partial Pressure of Oxygen assay (EPPO). Under these conditions, a complete absence of flavonoids does not affect dramatically seed longevity, as *tt4-8* seeds did not show more sensitivity than wild-type’s, and *tt5-1* seeds only show a weak sensitivity under EPPO. Moreover, *tt10-2* seeds do not present better longevity than wild-type’s, despite containing higher amounts of flavonoids. Depleting only quercetin-derived flavonols from the seeds does not affect longevity (as in *fls1-3*), and neither does it the absence of PAs, as in *tt3-4*, (or only partially, as in *ban-1*, whose longevity is reduced only a 25%) (Figure 1a, b). Remarkably, extreme sensitivity was observed in *tt7-4*, whose seeds presented a 60% lower germination than wild-type’s after CDT, and an 80% lower after the EPPO assay. This dramatic reduction was also observed in the *tt7-7* allele in Col-0 background (Supplemental Figure 1).

**Figure 1:**
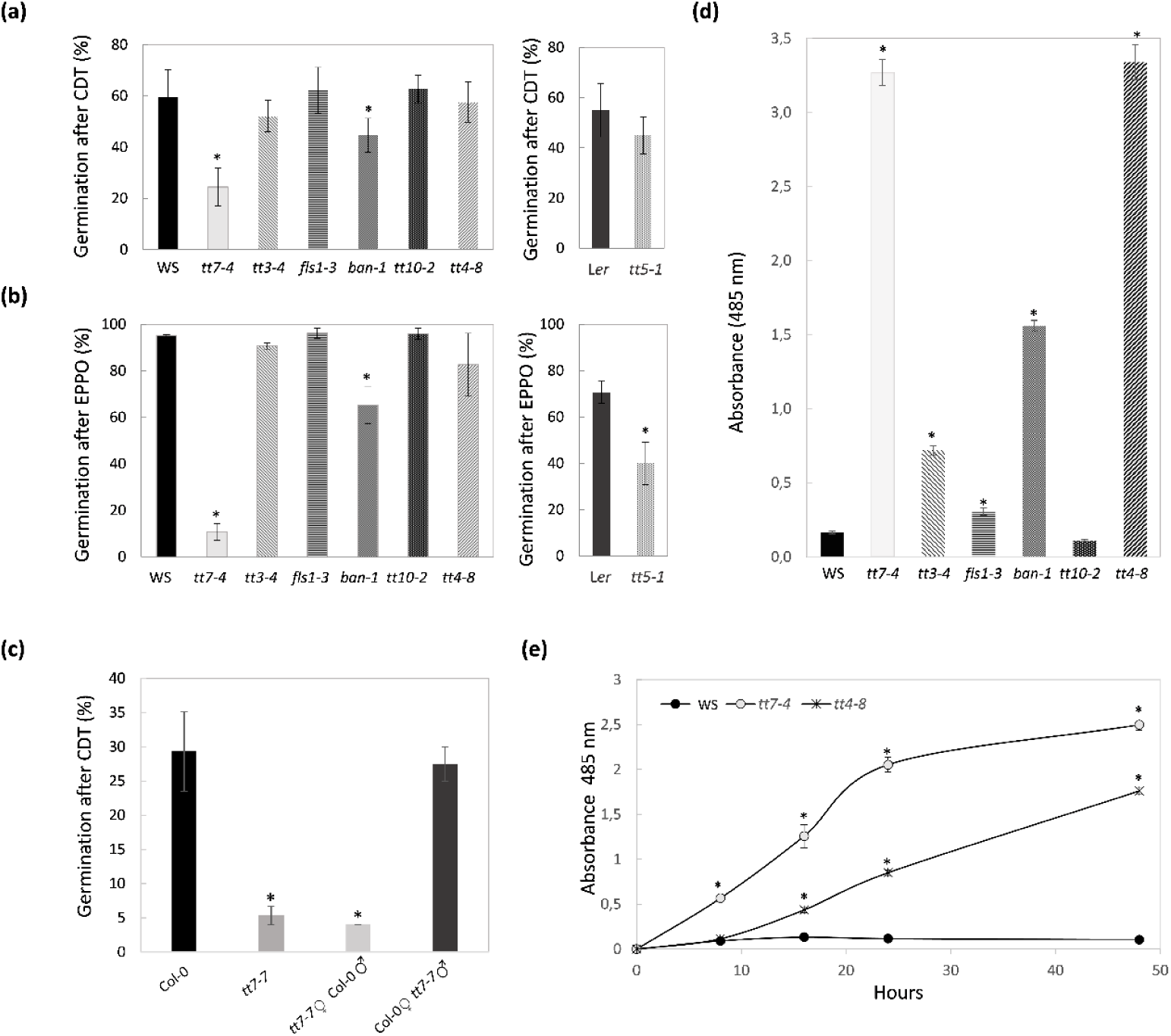
Altered seed longevity and permeability of *transparent testa* mutants and maternally inherited *tt7* reduced longevity. Seeds from 100%-germinating batches were incubated during 14 days at 75% RH in controlled-deterioration treatment (CDT) (a), and maintained 5 months at 5 bar O_2_ and 40% RH (Elevated Partial Pressure of Oxygen Assay, EPPO) (b) and percentage of germination was recorded after 5 days. c) Seeds from wild type (Col-0), *tt7-7* and F1 from reciprocal crosses were subjected to CDT (17 days at 75% RH), and percentage of germination was recorded after 7days. Bars represent the average and standard errors of three replicates with 30 seeds per line. WS, Wassilewskija. Ler, Landsberg *erecta*, Col-0, Columbia-0. d) Wild-type (WS) and mutant seeds were incubated during 3 days in 1% tetrazolium at 30°C and then formazan was extracted and quantified. e) Quantitative time course of formazan accumulation in WS, *tt7-4* and *tt4-8* seeds. Data (absorbance at 485 nM) are the mean and standard error of three biological replicates. Asterisks indicate significant differences with WT (P< 0.05) between samples in two-tailed t-Student test.

PAs are only accumulated in the seed coat, but flavonols are present both in maternal (seed coat) and in zygotic (embryo) tissues. To discern which of these two tissues harbour the seed longevity defect in *tt7*, we conducted reciprocal crosses. F1 seeds of wild-type plants that were pollinated with *tt7* pollen showed similar response as wild type after the ageing treatment. In contrast, F1 seeds of *tt7* plants pollinated with wild-type pollen retained the reduction in seed longevity after CDT observed for *tt7* (Figure 1c). These results, together with the fact that *tt7* fresh seeds do not present defects in germination (Supplemental Figure 2a), and that *tt7-7* embryos presented normal morphology and development (Grunewald et al, 2013, Supplemental Figure 2b-e), indicate that the maternal seed coat is responsible for the reduced seed longevity observed in *tt7* mutants after CDT.

Tetrazolium uptake is a proven method for analysing seed permeability in Arabidopsis seeds (Molina et al, 2008). Quantitative assays after 72h incubation revealed that the different *tt* mutants presented several degrees of staining (Figure 1d), indicating different rates of entry of tetrazolium through the seed coat. Control experiments ruled out the possibility that differences in staining are the result of defects in viability (Supplemental Figure 3a) or on the ability to reduce tetrazolium salts (Supplemental Figure 3b). In this assay, *tt3-4, tt4-8, tt7-4, ban-1* and *fls1-3* seeds were significantly more permeable to tetrazolium salts. In particular, a 1.8-fold increase in absorbance was observed in *fls1-3*, a 4.5-fold in *tt3-4*, and a 9.7-fold in *ban-1* seeds, compared to wild type. Intriguingly, *tt7-4* and *tt4-8* were by far the most permeable mutants, with increases of 20-fold. When the assay was performed after 48h of incubation, *tt7-4* showed higher staining than *tt4-8*, (Figure 1e), suggesting a faster entry of tetrazolium into the embryo. For *tt7-4, tt-3-4, fls1-3, tt10-2* and *ban-1*, higher permeability correlated well with reduced longevity (i.e., seeds were more sensitive to deterioration). Unlike these mutants, *tt4-8* seeds were not sensitive to CDT despite having highly permeable seed coats, probably indicating the existence of a compensatory effect in this mutant. Taken together, these results indicate that, among all mutants tested, *tt7* is the most sensitive to seed ageing. This sensitivity has a maternal origin and *tt7* seed coat is the most permeable among the mutants analysed here, suggesting a severe and distinctive defect in its seed coat.

### The outer integument is defective in the *tt7* mutant

Mucilage and lipid polyesters contribute to the physiological characteristics of the seed coat. We investigated the ability of flavonoid mutants to properly develop these two compounds using biochemical assays. When *Arabidopsis* wild-type seeds were hydrated, the seed coat mucilage swelled rapidly, and a pink-stained capsule was observed after ruthenium red staining. In contrast, in *tt7-4*, mucilage production was severely affected, being drastically reduced or absent (Figure 2a). After staining seeds with sudan red to visualize seed coat suberized cell walls, wild-type seeds showed the characteristic pink layer in the outer integument of the seed coat. This layer was not visible in *tt7-4*, suggesting that lipid polyester deposition is also compromised (Figure 2b). Chemical analysis of lipid polyesters composition in dry seeds revealed an overall ∼30% reduction in the *tt7-4* amounts of monomers released after depolymerization, compared to WT (Figure 2c inset). Most lipid monomers are significantly reduced in *tt7-4* compared to wild type controls, with the exception of three 22:0 straight chain suberin components (Figure 2c) and fatty acids with up to 22 carbons which showed higher accumulation in the mutant. The same results were observed in the *tt7-7* allele (Supplemental Figure 4a-b-c).

**Figure 2.**
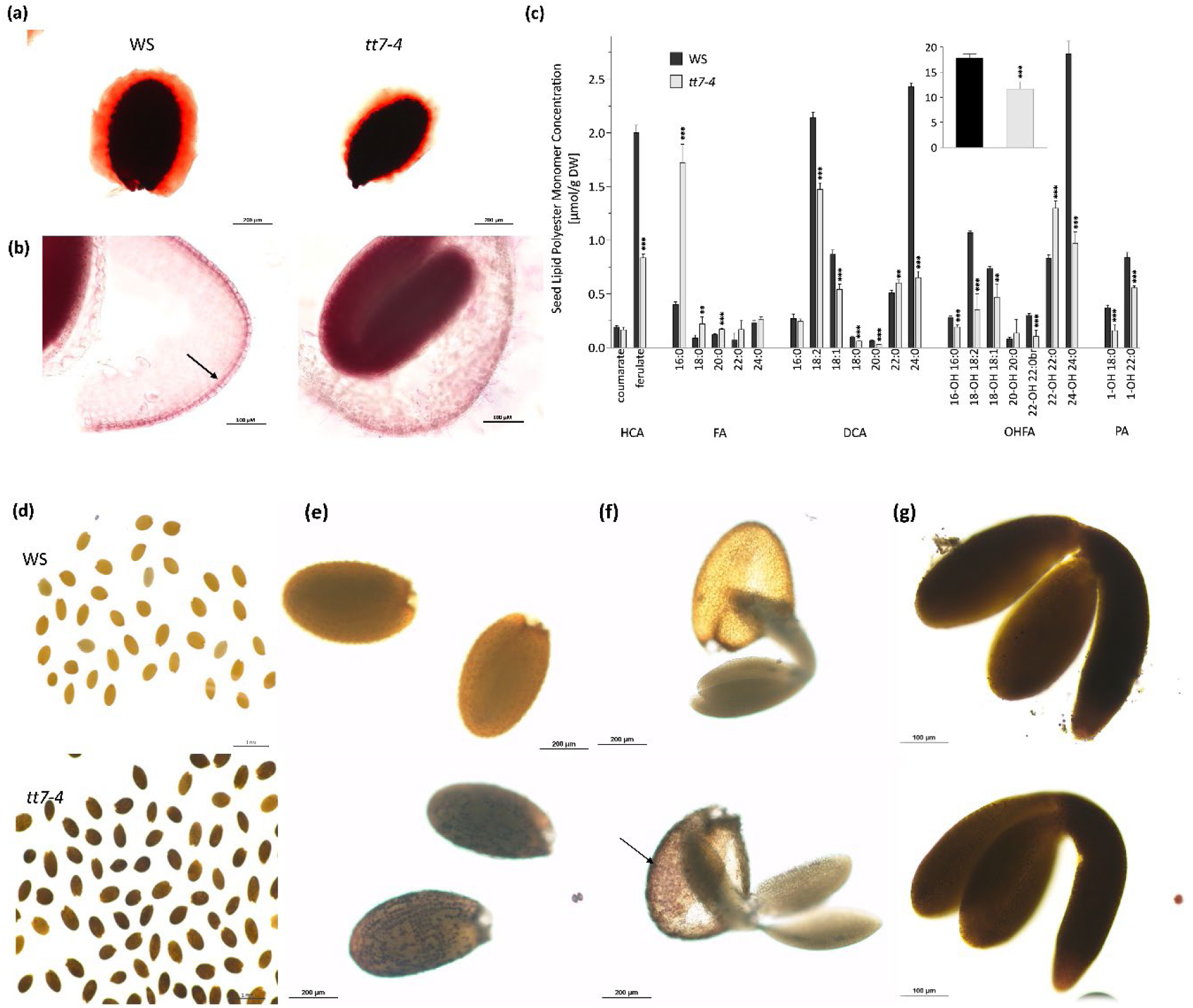
Seed coat development is affected in *tt7*. a) Representative images of WS and *tt7-4* seeds after staining with ruthenium red for 15 m. Scale bars: 200 µM. b) Representative images of WS and *tt7-4* seeds stained with Sudan Red 7B. Arrows indicate the palisade layer. Scale bars, 100 µM. c) Lipid polyester monomer composition of WS and *tt7-4* mutant seeds. Bars represent the mean content of each identified monomer and error bars represent SD. Asterisks indicate significant differences with WT (*, P< 0.05; **, P< 0.01; ***, P<0.001) between samples in two-tailed t-Student test. br: branched monomer; DCA: dicarboxylic acids; FA: fatty acids; HCA; Hydroxycinnamic acids; OHFA: hydroxyfatty acids; PA: primary fatty alcohols; WS, Wassilevskija. Inset: Total amount of lipid polyester monomers released after depolymerization of WS and *tt7-4* seeds. d) Image of WS and *tt7-4* dry seeds after iodine staining. Scale bar, 1mm. e) Amplified view of WS and *tt7-4* dry seeds after iodine staining. Scale bar, 200 µm. f) Representative images of WS and *tt7-4* seed coat and embryo from dry seeds stained with iodine. Only *tt7-4* seed coat (arrow) and not the embryo is stained. Scale bar, 200 µm. g) Representative images of WS and *tt7-4* developing embryos (12 DAP) stained with iodine showing that *tt7-4* embryos also synthetize starch at this stage. Scale bar, 100 µm. WS: Wassilevskija. DAP: Days after pollination.

These findings were confirmed by transmission electron microscopy (TEM) (Figure 3 a, b). In wild-type dry seeds, the columella was clearly observed in the most external layer of the outer integument. Beneath the epidermis lies the dark suberin-containing palisade layer (PL). In *tt7*, columella-like structures were identified, indicating that the outer cells secreted some mucilage that forced the protoplast towards the centre of the cell. However, this structure was only insinuated, being reminiscent of the initial stage of columella formation at the torpedo stage in wild-type, but without the characteristic starch granules at the center of the epidermal cells in this stage (Windsor et al, 2000). In the inner layer of the outer integument, the defined electron-dense layer of suberin of the palisade was not properly formed. Big granular structures of low electronic density, resembling starch granules present in wild-type seed coats at earlier stages, were highly abundant and distributed along the cells. The defects in *tt7* outer integument started at 9 DAP, when the developmental path of the two layers diverged. At 3 DAP, the five integumental layers were distinguished in *tt7* seed coats (Supplemental Figure 5). At 7 DAP, the highly vacuolated large cells of the two external layers were observed in wild type, together with the endothelium beneath them, with the electron-dense tannin vacuole covering the whole cytoplasm. This same structure was present in *tt7*, being the only distinct feature the appearance of large round globules in the cytoplasm of the endothelium cells, that could contain the epiafzelechin-derived PAs characteristic of *tt7* (Figure 3c). Similar to wild type, the *tt7* epidermis did not contain granules and produces mucilage at 9 DAP, with cytoplasm forced to generate pin-like structures perpendicular to the seed surface. In the inner layer, however, starch-resembling granules continued to accumulate, pushing the vacuole towards the inner region of the cell. The cell walls were not reinforced, which explains the less-defined structure of this layer (Supplemental Figure 6). At 16 DAP, *tt7* columella is clearly defective, and the inner cells still contained granules, that remained present until maturity.

**Figure 3:**
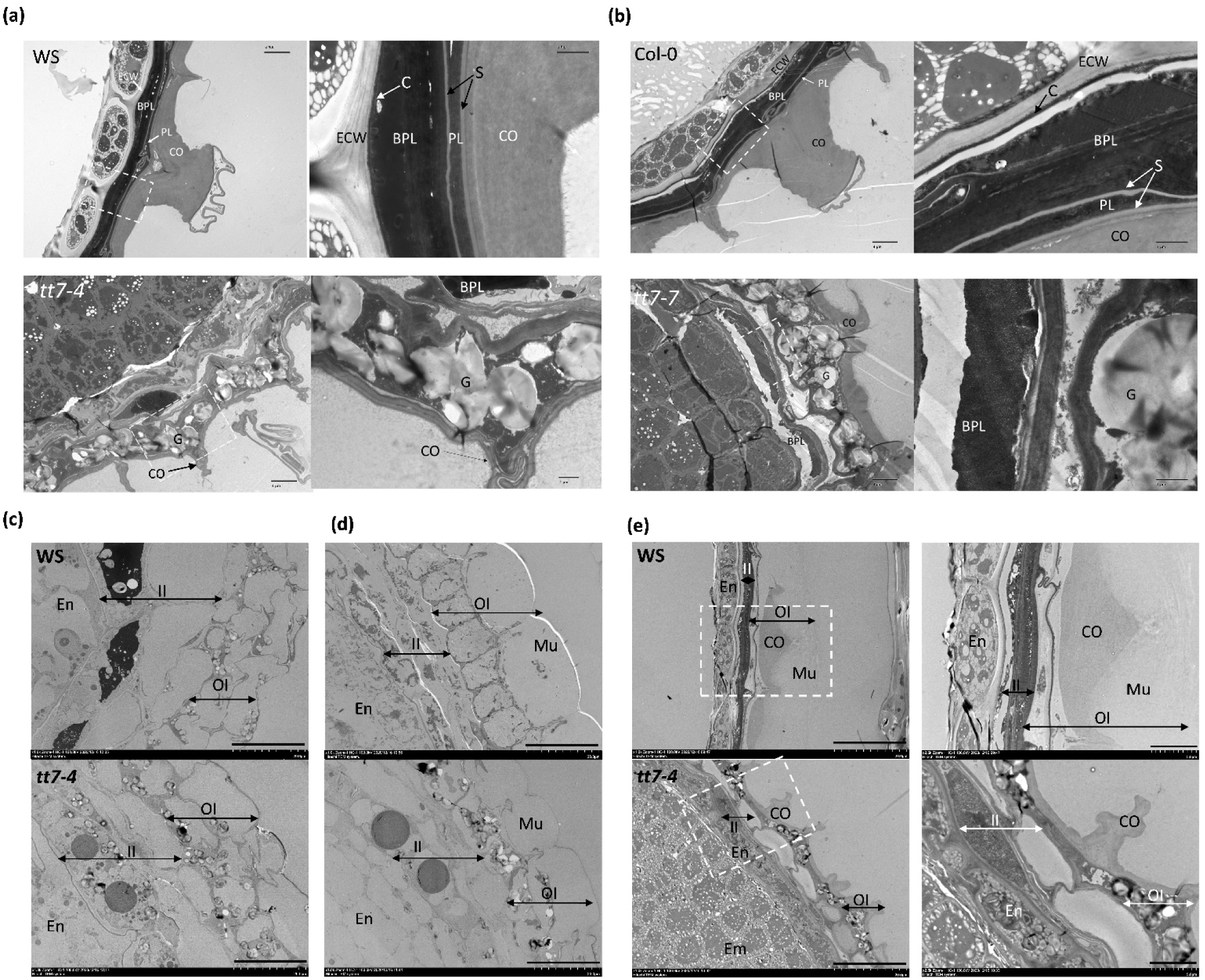
TEM images of a representative seed coat of wild type and *tt7* seeds. **a) WS** and *tt7-4* dry seeds. b) Col-0 and *tt7-7* dry seeds. White boxes indicate the area amplified in right panels. Left panels scale bars, 4 µmM; Right panels scale bars, 1 µm; Representative TEM images of seed coats of WS (above) and *tt7-4* mutant (below) at 7 days (c), 9 days (d) and 16 days (e) after pollination. White boxes in e) indicate the zoomed areas in right panels. Scale bars in c), d) and e) left panel: 20 µm; Scale bars in e) right panel, 5 µm; En: Endosperm; ECW: Endosperm cell wall; BPL: Brown pigment layer; PL: Palisade layer; CO: Columella, G: Granules; II: Inner integument; OI: Outer integument; Mu: Mucilage, CO: Columella; WS: Wassilevskija; Col-0: Columbia 0.

Iodine staining was performed to confirm that the low electronic density granules observed by TEM in the inner layer of the outer integument of *tt7* dry seeds contained starch. As expected, wild-type dry seeds were not stained, as starch accumulated during seed coat development has been properly degraded at maturity. However, an intense blue staining was observed in *tt7* dry seeds (Figure 2d-e and Supplemental Figure 7). Interestingly, this starch accumulation effect was seed coat-specific, as embryo (Figure 2f-g; Andriotis et al, 2010) and leaves starch (Supplemental Figure 8) was properly synthetized and degraded. In sum, these results indicate that differentiation of the seed coat outer integument (specifically seed coat lipid polyesters synthesis and seed coat starch metabolism) is altered in *tt7* mutants.

### Kaempferol accumulation is causing the seed coat and longevity defects of *tt7*

We next investigated whether the defects observed in *tt7* seed coats were shared by other flavonoid mutants. *tt3-4, tt4-8, tt10-2*, and *fls1-3* seeds were able to produce a layer of mucilage of similar thickness than wild type. In *ban-1* and *tt5-1*, the amount of mucilage was reduced, but it is still observed. The lipid polyester layer in all mutants was also similar to wild type (Figure 4a). Seed coat starch accumulation in *tt7-4* was not shared with the other mutants (Supplementary Figure 9). These findings were confirmed by TEM observation of *tt4-8, fl1-3* and *ban-1* seed coat sections at maturity (Figure 4b). In these mutants, the ultrastructure of the seed coat outer integument was similar to that of WT seeds. Therefore, the altered flavonoid pattern of *tt7* seems to be causing its distinctive phenotypes. A flavonoid deficient *tt7-4 tt4-8* double mutant was generated to confirm this hypothesis. The *tt7* defects in lipid polyesters and starch accumulation were eliminated in the *tt7-4 tt4-8* seeds (Figure 5a, b). Similarly, the sensitivity to ageing of *tt7-4* seeds was abolished in *tt7-4 tt4-8*, indicating that *tt7* defects are flavonoid-dependent (Figure 5c).

**Figure 4:**
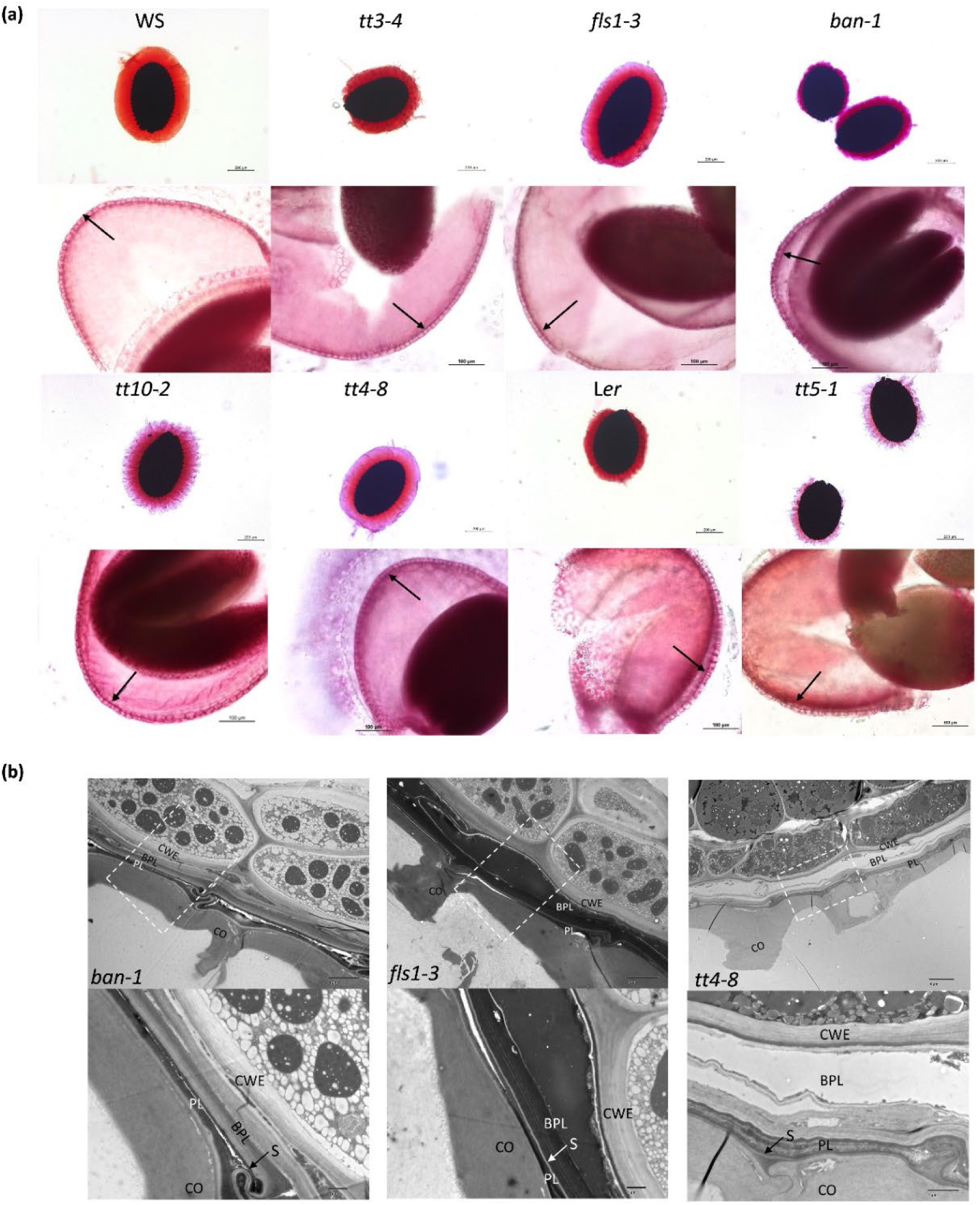
*tt7* seed coat defects are not shared with other *tt* mutants. a) Ruthenium red (above) and Sudan Red7 (below) staining of different *tt* mutants showing mucilage and polyester layers. Scale bars 200 and 100 µm, respectively. Arrows indicate palisade layer. b) TEM images of a representative seed coat of *ban1, fls1-3* and *tt4-8* mutants. White boxes indicate the area amplified in panels below. Left panels scale bars, 4 µmM; Right panels scale bars, 1 µm; ECW: Endosperm cell wall; BPL: Brown pigment layer; PL: Palisade layer; CO: columnella. WS: Wassilevskija.

**Figure 5.**
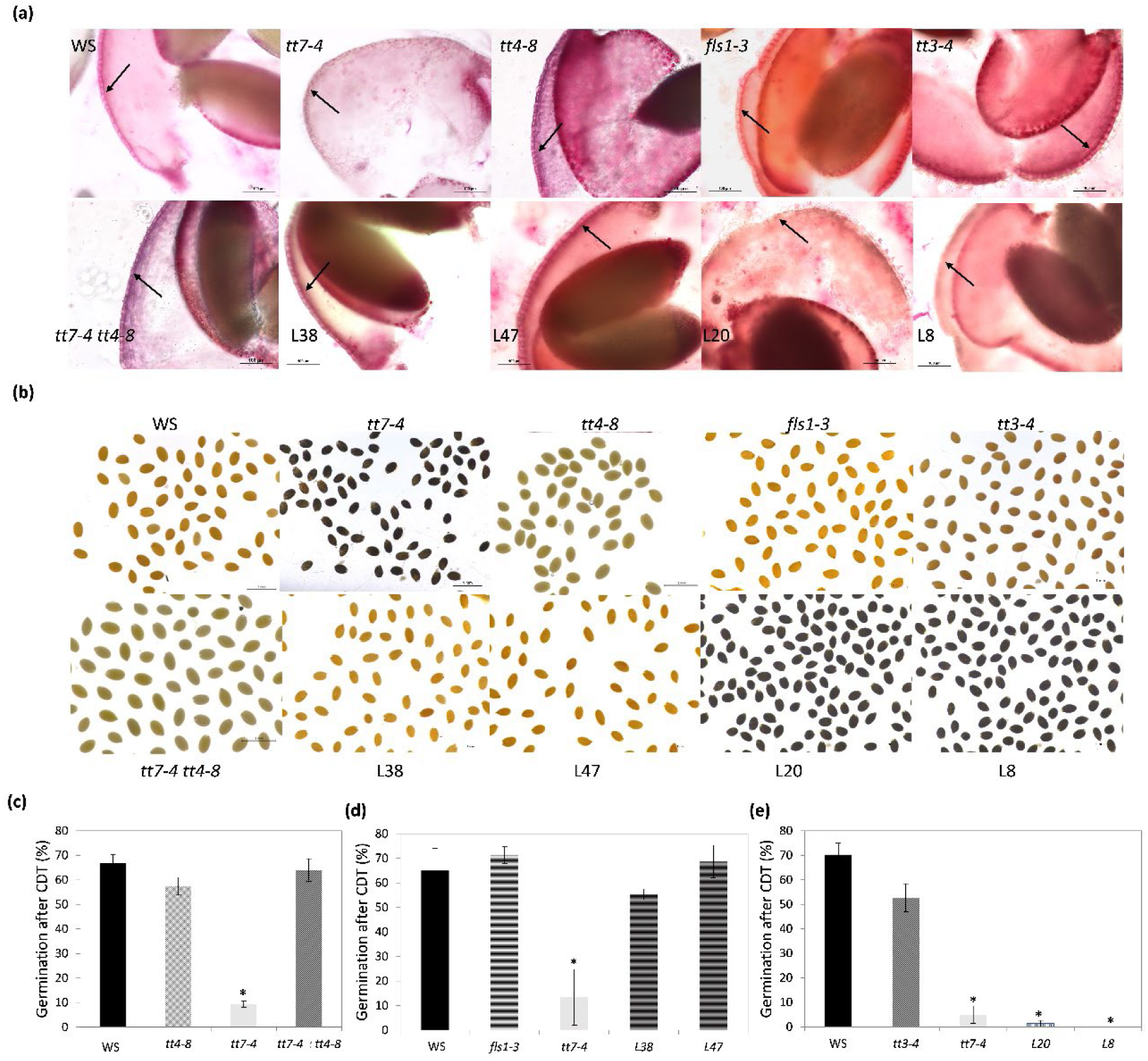
*tt7* seed coat defects are due to over-accumulation of a kaempferol-derived compound. Representative image of WS, *tt7-4, tt4-8, fls1-3, tt3-4, tt7-4 tt4-8*, two lines (L38 and L47) obtained by directed mutation of TT7 gene in *fls1-3* mutant background, and two lines (L20 and L8) obtained by directed mutation of TT7 gene in *tt3-4* mutant background. a) Sudan Red7 staining, scale bars 100 µm. Arrows indicate palisade layer. b) Iodine staining. Scale bar, 1mm. c-e) Percentage of germination after CDT. Seeds were incubated during 14 days at 75%. Percentage of germination was recorded after 5 days. Bars represent the average and standard errors of three replicates with 30 seeds per line. WS, Wassilewskija. Asterisks indicate significant differences with WT (P< 0.05) between samples in two-tailed t-Student test. CDT: Controlled Deterioration Treatment.

Two features are characteristic of the *tt7* flavonoid profile: a dramatic increase of kaempferol-derived flavonols, and an altered composition of PAs (Routaboul et al, 2016). To discern which of these two groups of flavonoids is responsible of the *tt7* phenotype, *TT7* was chosen as target to generate CRISPR-edited plants in *tt3-4* or *fls1-3* mutant backgrounds. A *tt3 tt7* double mutant would accumulate kaempferol, but would not synthesize PAs, whereas *fls1 tt7* would not synthesize kaempferol but would accumulate more PAs. To generate these lines, we used the promoter of the egg cell-specific EC1.2 gene to drive the expression of Cas9, and two guide RNAs to create an internal deletion in the target genes, leading to homozygous knock-out mutants in the T1 generation (Wang et al, 2015 Supplemental Methods). Seeds from T1 *fls1-3* or *tt3-4* plants, homozygous for CRISPR-edited *tt7* (cr), where biochemically assayed for seed coat components (lipids and starch), and seed longevity. As shown in Figure 5 a,b and Supplemental Figure 10, *fls1-3 tt7cr* seeds (L38 and 47) lost the *tt7* phenotypes and recover wild-type appearance. In addition, they lost the sensitivity to ageing observed in *tt7-4* (Figure 5d). Conversely, *tt3-4 tt7cr* seeds (L20 and L8) retain the structural *tt7* defects in the outer integument (Figure 5a,b; Supplemental Figure 10). Moreover, they retain the reduced seed germination after CDT characteristic of *tt7-4* (Figure 5e). Taken together, these results indicate that the defects observed in *tt7* seed coat are due to the excess accumulation of kaempferol or its derivatives in this tissue.

### Starch degradation in the seed coat is needed for seed longevity

We recently demonstrated the role of suberin in the regulation of seed permeability and longevity (Renard et al, 2020, Renard et al, 2021). According to this, the sensitivity to ageing of *tt7* seeds can be partially explained by its defective suberin layer. To determine if starch accumulation could also be influencing seed longevity, a genetic approach using mutants accumulating starch in the seed coat was employed. Moderate starch accumulation in mature seeds of *sex1* and *sex4* is described (Andriotis et al, 2010). In leaves, the most severe starch phenotype appears in the triple mutant *amy3 isa3 lda*, which completely prevents starch breakdown (Streb et al, 2012). Despite the wilt-type pattern of seed starch accumulation of the single *amy3* and *isa3* mutants (Andriotis et al, 2010), this triple mutant strongly accumulates starch in dry seeds, especially in the seed coat (Supplemental Figure 11a, c). Longevity effects were not observed for *sex1-1, sex4-3 isa3-2* and *sex4-3 amy3-2* after our CDT assay, suggesting that the defects of *sex1* or *sex-4* are not sufficient to compromise longevity. Interestingly, *amy3 isa3 lda* was very sensitive to ageing, with only 38.8% germination versus 72.7% in wild type, suggesting that when starch breakdown is severely blocked in the seed coat, longevity is affected (Figure 6). The *amy3 isa3 lda* mutant does not present a dramatic increase in seed permeability (Supplemental Figure 11b), confirming that seed coat starch is not used as a reserve for lipid biosynthesis, representing a novel component involved in seed longevity independent of this property.

**Figure 6:**
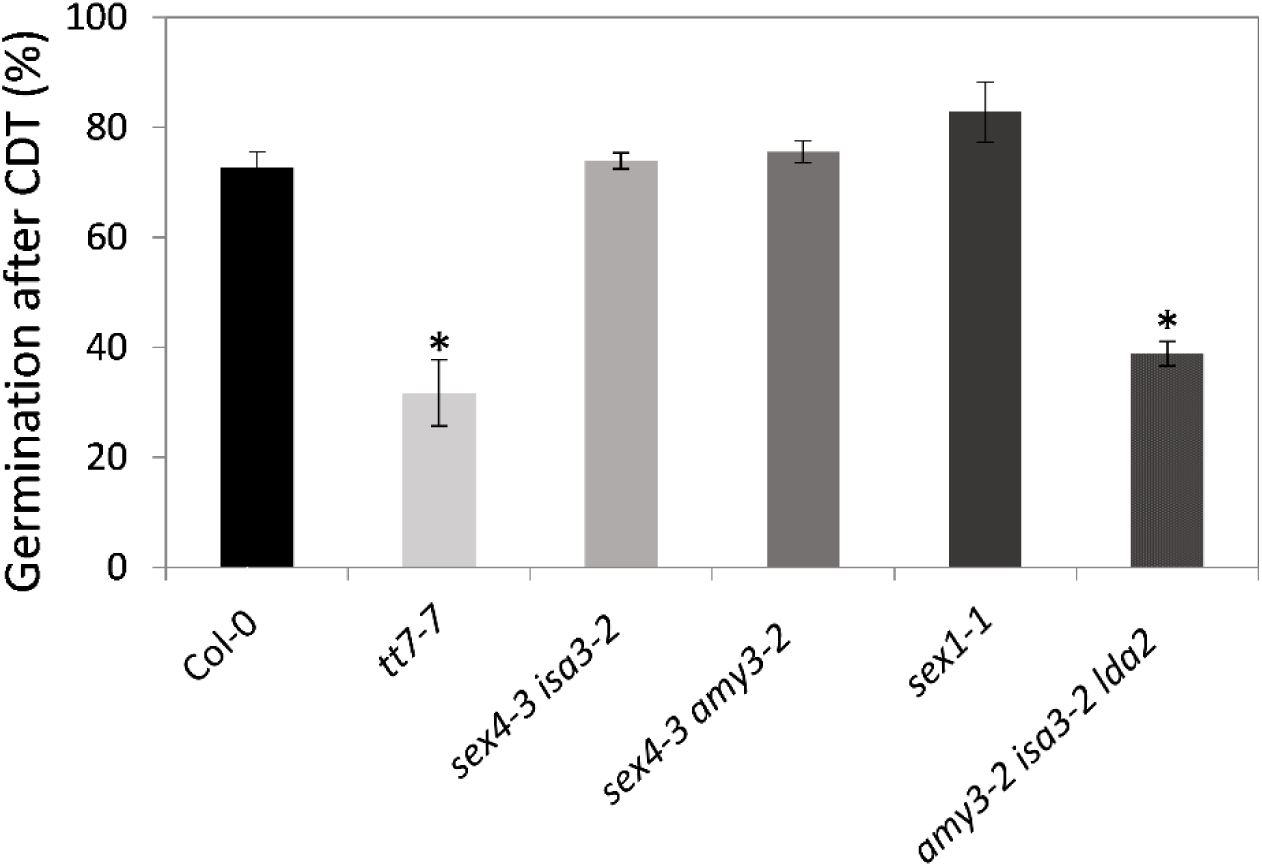
Seed coat starch accumulation affects seed longevity. Percentage of germination of Col-0, *tt7-7, sex4-3 isa3-2, sex4-3 amy3-2, sex1-1* ans *amy3-2 isa3-2 lda2* mutant seeds after CDT. Seeds were incubated during 14 days at 75% RH and sown on MS plates. Percentage of germination was recorded after 5 days. Bars represent the average and standard errors of three replicates with 30 seeds per line. Col-0, Columbia 0. Asterisks indicate significant differences with WT (P< 0.05) in two-tailed t-Student test. CDT: Controlled Deterioration Treatment.

### Expression of starch synthesis and degradation genes is higher in *tt7* seeds

Seed coat starch begins to degrade at 9 DAP, and *tt7* defects extends up to maturity, when starch granules are still observed in the outer integument. To discern whether the increased content of starch observed in *tt7* is due to a repression of starch-degradation genes, expression was assayed in seeds of 10-14 DAP. As wild-type and *tt7* embryos behave similarly regarding embryo seed starch synthesis and degradation (Figure 2f-g), differences in expression are attributed to seed-coat starch. Unexpectedly, expression of most of the starch-degrading genes analysed was increased in *tt7* seeds (Figure 7a). This include the chloroplastic *SEX1* and *AMY3* genes, but higher differences were found in the genes encoding the chloroplastic β-AMYLASES BAM1 and BAM3 and the nonenzymatic regulator BAM9, as well as in the cytosolic β-AMYLASE BAM5, GLUCAN-WATER-DIKINASE GWD2 and DISPROPORTIONATING ENZYME DPE2. Accumulation of starch granules depends on the rates of starch synthesis and degradation. Therefore, we analysed also the expression of starch-synthesis genes. Intriguingly, expression of the SUCROSE SYNTHASE SS4, the ADP-GLUCOSE PYROPHOSPHORYLASE ADG1 and the PHOSPHOGLUCOMUTASE PGM1 was also found increased in *tt7* (Figure 7b). These findings indicate that starch accumulation in *tt7* seed coat is not due to a transcriptional downregulation of starch-degrading enzymes. In addition, it suggests an enhanced rate of starch biosynthesis.

**Figure 7.**
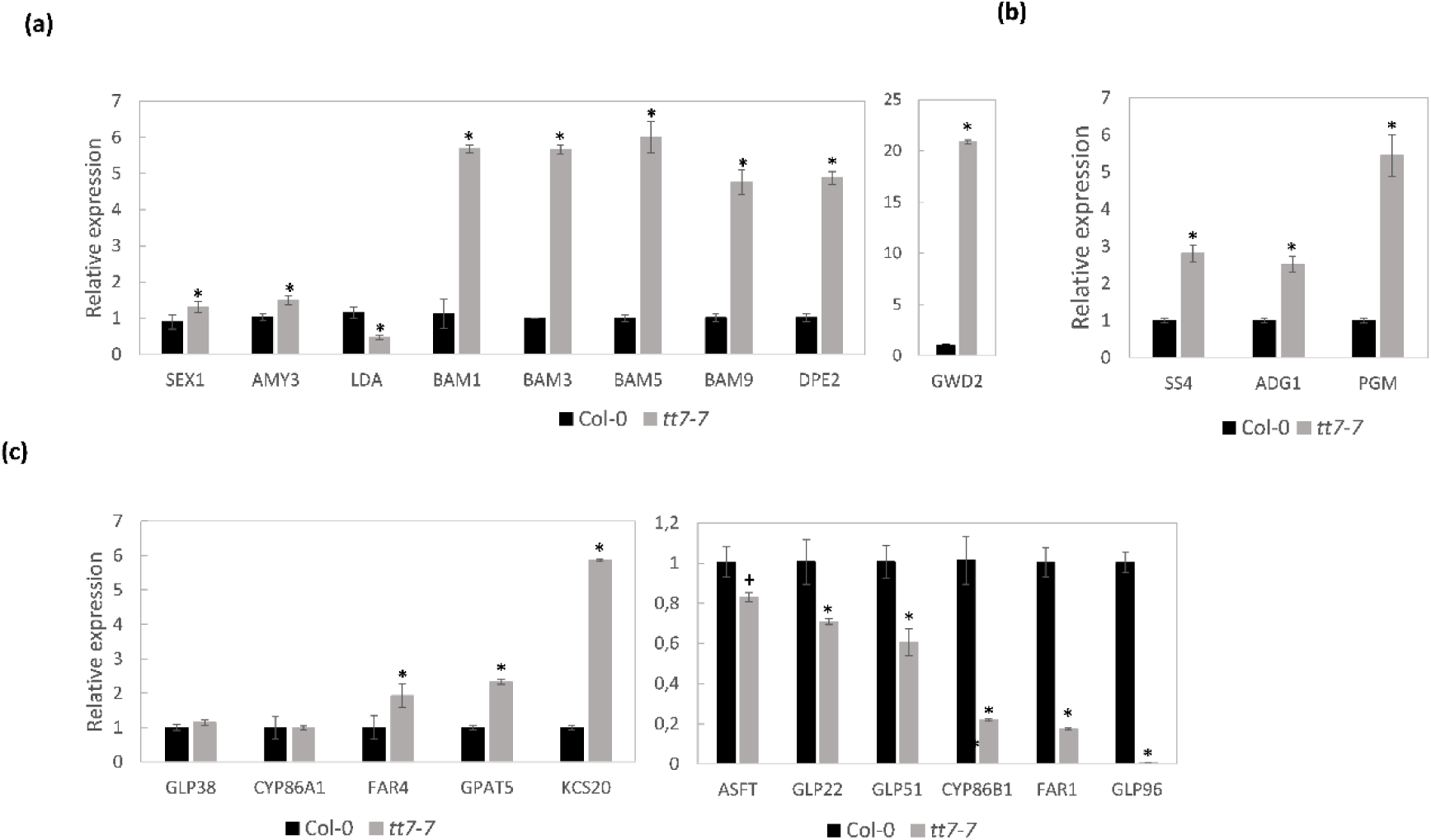
Expression of starch metabolism and suberine biosynthesis genes on *tt7*. Expression levels of starch degradation (a), starch biosynthesis (b) and suberin biosynthesis genes (c) was determined by real-time quantitative PCR in developing seeds (10-14 DAP) of Col-0 and *tt7-7*. Data are the mean of three replicates. Expression for every gene in Col-0 is set to 1 in the y-axis and relative expression in *tt7-7* is shown. Significantly differing from wild type at P<0.05 (*) or 0,1 (+) using student’s t-test. Col-0=Columbia 0.

### Expression of suberin biosynthesis genes is lower in *tt7* seeds

To discern whether suberin synthesis in *tt7* is repressed at the transcriptional level, the expression of genes involved in suberin biosynthesis was assayed in seeds of 10-14 DAP, when this process is taking place in the outer integument. As shown in Figure 7c, the *FATTY ACID REDUCTASE (FAR1)*, the *VERY LONG CHAIN FATTY ACID HYDROXYLASE (CYP86B1)* and the *FERULOYL-COENZYME A TRANSFERASE (ASFT)* genes were downregulated in *tt7-7* seeds, explaining the reduction of suberin monomers observed in Supplemental Figure 4c. Three GDSL-lipases *GELP22, GELP51* and *GELP96* genes required for suberin polymerization (Ursache et al, 2021) were also found downregulated in *tt7*. Fatty acid reductase FAR1 and *FATTY ACID OMEGA-HYDROXYLASE* (*CYP86A1*) expression was not altered in *tt7* seeds, and *GLYCEROL 3-PHOSPHATE ACYLTRANSFERSASE5* (*GPAT5*) and *FATTY ACID ELONGASE* 20 (*KCS20*) were upregulated. These data indicate that the altered content of suberin monomers in *tt7* seeds can be due to the transcriptional downregulation of key biosynthetic genes, and suggest also a defect in lipid polymerisation.

### Auxins mediate expression of suberin and starch biosynthesis genes, but auxin localization in *tt7* is similar to wild type

Flavonols modulate auxin transport (Peer and Murphy, 2007), so our first hypothesis was that auxin signalling could be compromised in *tt7* mutants. Auxins could be playing a role in suberin and starch biosynthesis, as expression of SS4, ADG1 and PGM1 is activated by auxins (Zhang et al, 2019), and expression of the GDSL-lipases is repressed by auxins (Ursache et al, 2021). Moreover, as shown in Supplemental Figure 12, the expression of the *tt7*-repressed genes *FAR1* and *ASFT* were significantly reduced upon auxin treatment, suggesting that delocalization of auxins could be causing suberin biosynthesis repression and also starch biosynthesis activation in the *tt7* seed coat. However, auxin presence, as revealed by the AUXIN RESPONSE5 promoter:GFP (DR5:GFP) marker line (Weijers et al, 2006) was found similar in wild type and *tt7* seeds, being detected at the chalazal end of the seed coat up to 11 DAP (and not in the outer integument) (Supplemental Figure 13). Using this tool, auxins do not seem delocalized in *tt7* seed coat, and their involvement in suberin and starch biosynthesis in the seed coat requires further investigations.

### Global expression analysis reveals that cell trafficking and cell communication processes are altered in *tt7* seeds

To provide more clues about possible pathways affected by TT7 activity, transcriptome profiling was performed at 8 DAP (the day after the peak of TT7 expression during seed development, Winter et al, 2007) on Col-0 and *tt7-7* mutant seeds. A total of 4289 genes showed significant differential expression (DEGs) between the *tt7* and WT (Supplemental Table 1). Pruned GO term analysis among DEGs in the mutant showed enrichment of 313 functional categories in the Biological Process, and 118 in the Molecular Function Gene Ontology classification, indicating that disruption of TT7 activity provokes a dramatic readjustment of cell biology. Among the categories enriched in the 2284 DEGs *tt7* upregulated genes, shown in Supplemental Table 2 (the top 20 non-redundant terms are shown in Figure 8a-b), it is remarkable the appearance of cytoskeleton-related categories, such as actin filament-based movement (GO:0030048), which includes the concerted upregulation of myosins, and phosphatidylinositol-related processes (GO:0046488, GO:0052866, GO:0016307), including the upregulation of phosphatidylinositol-kinases and phosphatases. Cell communication-related categories, such as signalling (GO:0023052), movement of cell or subcellular component (GO:0006928) and cell communication (GO:0007154) include other the coordinated upregulation of kinesins, as well as some calcium sensors, Rho GTPases and other kinases. Finally, the category (1->3)-beta-D-glucan biosynthetic process (GO:0006075) was enriched by the concerted upregulation of eight isoforms of callose synthase genes, involved in plasmodesmata regulation. Detailed differential expression data of genes belonging to these categories is shown in Supplemental Table 1. Among the 2005 DEGs that were downregulated in *tt7*, it is remarkable the enrichment of categories such as plant-type cell wall organization or biogenesis (GO:0071669), including the downregulation of expansins and cellulose synthases, among other cell wall enzymes, lipid localization (GO:0010876), including the downregulation of lipid transfer proteins, and lipid metabolism (GO:0006629), with the downregulation of diverse lipid metabolism enzymes (Figure 8c-d). Thus, the transcriptomic results indicate that *tt7* mutants have severe defects in cellular components such as the cytoskeleton or plasmodesmata, as well as disfunction of molecular switches involved in cell signalling, organelle trafficking and lipid biology.

**Figure 8.**
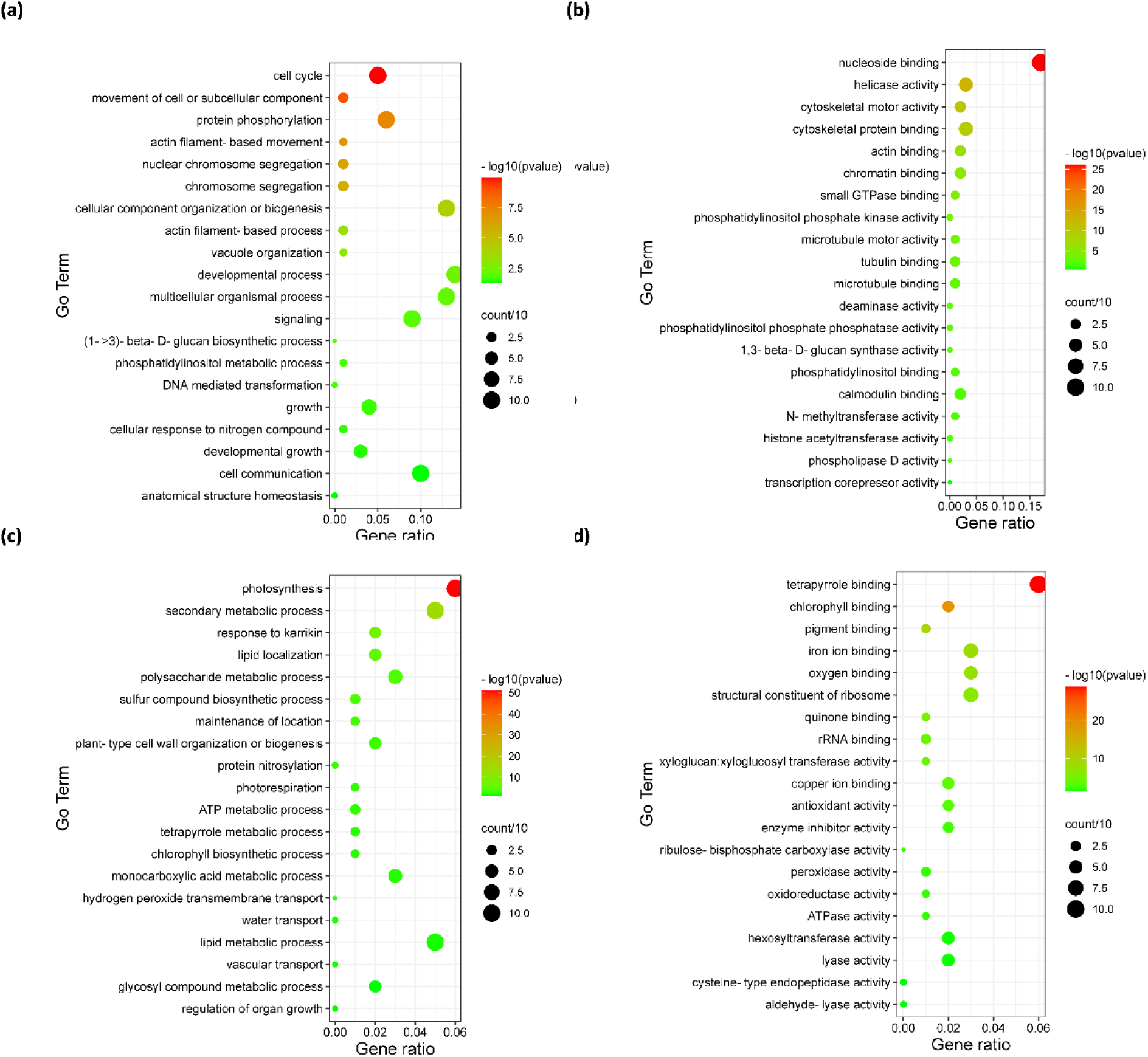
GO categories enriched on *tt7* differentially-expressed genes. Top-20 non-redundant enriched GO categories in *tt7* upregulated (upper panel) and downregulated (lower panel) genes. a) and c) refer to Biological Process Terms, b and d) refer to Molecular Function Terms. GO enrichments were obtained using AgrigoV.2 and filtered using REVIGO to remove semantically redundant terms. Terms were pruned by REVIGO frequency (terms with frequency >10% were removed) and ranked by dispensability. Full lists can be found in Supplemental Table 2. a) Biological Process GO categories, b) Molecular Function GO categories. Gene ratio: Ratio of upregulated genes in a given category divided by total number of genes in this category. Counts: Number of upregulated genes in a given category. -log p (p-value in log scale after false discovery rate correction).

## DISCUSSION

Despite being one of the best-studied secondary metabolite pathways in plants, the function of flavonoids in the seed coat is not totally understood. Considered important for seed longevity since Debeaujon et al (2000) showed that a set of mutants defective in flavonoid biosynthesis presented increased seed deterioration upon ageing, it is still uncertain what is the link between the defects in these mutants and seed longevity, as well as how essential flavonoids are for this trait. Their antioxidant properties were invoked to explain this longevity effect, although flavonoid contribution to the integrated antioxidant system of plants is under debate (Hernández et al, 2009). Arabidopsis seeds accumulate flavonols and proanthocyanidins (PAs) (Lepiniec et al, 2006). Given their seed-coat specificity and that the *transparent testa* phenotype refers to the lack of brown colour provided by these compounds, PAs received more attention than seed coat flavonols as candidates to be the main flavonoids involved in seed protection. Without neglecting a contribution to sustain oxidative damage from outside during storage, this does not explain why mutant seeds completely deployed of PAs but maintaining its content in flavonols (*tt3-4*), or with increases of PA amounts (*fls1-3, tt10-2*) or changes in PA composition (*tt7-4 fls1cr*) present a similar rate of deterioration than wild type seeds after EPPO treatment, which mimic oxidative conditions (Groot et al, 2012). Even the complete absence of flavonoids (including PAs, *tt4-8*) does not impact seed viability. Results by Debeaujon et al, 2000 show subtle differences with this study regarding the relative sensitivity to some of the mutants, likely due to differences in the accessions used, which are known to present differences in seed longevity (Renard et al 2020b) and could confuse interpretation by overlapping two variables. In both studies, however, TT4 lack of function does not affect seed longevity (as also reported by Clerkx et al, 2004), suggesting that the intrinsic presence of PAs (or flavonols) in the seed coat, and hence their antioxidant potential, is not the main player determining viability. Another role for PAs in seed longevity argues to their structural function, binding to cell walls and providing physical protection (Pourcel et al, 2006). PAs are structural components of the endosperm cuticle, a layer compromised in several *tt* mutants with enhanced seed permeability (Loubéry et al, 2018). The low longevity of the flavonoid mutants found by Debeaujon et al, 2000 would be then attributed to the lack of PAs and higher permeability of their cuticular layer. Outer integument suberin also contributes to impermeability, and this layer is synthesized later in development, when seed longevity is being acquired (Riguetti et al, 2015). It covers the entire seed coat, a requisite to avoid entrance of molecules from outside (Molina et al, 2008). Mutant seeds affected in suberin synthesis have increased seed permeability and longevity and thickening this layer increase seed longevity in several species (Renard et al, 2021). Among all the mutants assayed in this study, *tt7* presents the most dramatic reduction of longevity, and in contrast with others, it also shows distinctive defects in the outer integument histology and cell wall suberization. *tt7* is also the mutant showing the highest permeability, which would explain its extremely low viability. Interestingly, our study revealed that seeds severely defective in seed coat starch degradation also present reduced viability, adding a new component contributing to this trait independent of seed permeability. Digestion of starch granules in outer integument is needed for proper cell differentiation (Andriotis et al, 2010), and contributes to the structural formation of the seed surface via reinforcement of cell walls (Kim et al, 2005), which protect the seed from external damage. These findings suggest that, without neglecting the contribution of the endothelium cuticle to seed permeability (Loubéry et al, 2018), the most essential barriers determining seed longevity are located in the outer integument, and pinpoints flavonols as regulators of its differentiation. It is plausible that the longevity reduction of other mutants defective in outer integument development, such as *ap2* (Western et al, 2001), could also be caused by an altered homeostasis of flavonols. In this regard, a recent report shows that the miR172-mediated repression of AP2 alters flavonoid biosynthesis in apple (Ding et al, 2022). Curiously, defects in lipid polyesters (suberin and cutin) have been also observed in the *transparent testa* mutant *tt15* (DeBolt et al, 2009; Loubéry et al, 2018). Although its flavonoid profile is altered, it does not over-accumulate kaempferol derivatives (Routaboul et al, 2012), indicating that more than one flavonoid subgroup could be interfering with suberin biosynthesis. The defects in *tt15*, however, are different from those observed in *tt7*: the inability to degrade starch is not reported for *tt15*, although its cuticle is affected (Loubéry et al, 2018).

The defects of *tt7* seeds are restricted to the seed coat, pointing to a seed coat-specific kaempferol derivative interfering with outer integument normal development. Total seed kaempferol content was raised by 63% in *tt7-4* (Routaboul et al, 2006), and the mutation led to the accumulation of kaempferol aglycone and kaempferol-glycoside derivatives, including a new kaempferol-rhamnoside-glycoside, not found in other *tt* mutants. The nature of the flavonol causing the seed coat defects in *tt7* is currently under investigation. Interestingly, alterations in kaempferol derivates profile have a tremendous impact in plant development. The modified flavonol glycosylation profile in the *rol1-2* mutant (with increases in kaempferol-3-O-rhamnose, kaempferol-3-O-glucose and another unknown kaempferol-derived compound) provokes hyponastic growth, aberrant morphology in cotyledons and defective trichome formation (Ringli et al, 2008). Similarly, mutations in the UGT78D2 gene, encoding a 3-O-glucosyltransferase, also present an altered kaempferol glycoside pattern and develop a dwarfing shoot phenotype and loss of apical dominance (Yin et al, 2014). Here, the active compound responsible of the *ugt78d2* phenotype was attributed to kaempferol 3-O-rhamnoside-7-O-rhamnoside. Assaying seed development and longevity in these and others flavonol glycosylation mutants or kaempferol over-accumulators (Sotelo-Silveira et al, 2013) may indicate whether *tt7* phenotypes can be traced back to a single kaempferol compound.

Evidence has accumulated implicating flavonols in plant growth and auxin transport modulation (Peer et al, 2004; Kuhn et al, 2011). Auxins also drive roles during seed longevity acquisition in the embryo (Pellizzaro et al 2019). Whether these processes are modulated by flavonoids is unknown, but the fact that TT7 is the only flavonoid biosynthetic gene found in the coexpression network during longevity acquisition in *Medicago* and *Arabidopsis* (Riguetti et al, 2015) points in this direction. Downregulation in *tt7* of auxin-regulated suberine biosynthetic genes suggest that suberization of the seed coat could be also mediated by auxins, and that the modified flavonoid profile in *tt7* disrupts this regulation. The altered expression of phosphatidylinositol kinases (PIPKs) and cytoskeleton-related genes in *tt7* seeds are additional evidence to connect flavonols with auxin transport in seeds, given their involvement in this process (Ischebeck et al, 2013; Ojangu et al, 2018; Abu-Abied et al, 2018; Song et al, 2021). Besides, auxin transport is repressed in *tt7* seedlings, and auxin efflux carriers PIN1 and PIN4 are delocalized (Peer et al, 2004). Therefore, finding a similar auxin localization (at least up to 11 DAP) in wild-type and *tt7* seeds was unexpected. Whether auxins are involved in these processes and whether they are playing a role in *tt7* defects requires further investigations.

Kaempferols inhibit activity of PIPKs (Kong et al, 2011) and Rho GTPases (Sharma et al, 2019), and are able to bind actin (Böhl et al, 2007). The link between these three proteins in cytoskeleton function is well-established (Gu et al, 2005; Davis et al, 2007; Spiering & Hodgson, 2011; Croisé et al, 2014). The transcriptomic profile of *tt7* seeds suggest disfunction of these proteins and insinuates that the cytoskeleton is not functioning properly in this mutant. Consequently, vesicle trafficking would be compromised, affecting seed coat cell-wall synthesis and remodelling (Krishnamoorthy et al, 2014) and localization of membrane-bound suberine transporters or lipid transfer proteins involved in suberine formation (Deeken et al, 2016), which were found deregulated in *tt7* seeds. Actin cytoskeleton (Su et al, 2010; Chen et al, 2010) and transient callose deposition (Vatén et al, 2011; Wu et al, 2018; Cheval et al, 2020) are also mechanisms to control plasmodesmata aperture. In *tt7* seeds, the coordinated upregulation of callose-synthase genes indicates that cell-cell communication could be affected. Upregulation of callose synthases could mean that plasmodesmata channels are closed and that symplastic communication is reduced in *tt7* seeds, which would alter the traffic of mobile signalling molecules for normal development. Interestingly, the *cad1-3* mutant, deficient in the *PHYTOCHELATIN SYNTHASE* gene, has increased levels of kaempferol 3,7 dirhamnoside and is also impaired in callose deposition in the plasmodesmata after Cd exposure (De Benedictis et al, 2018). These results open the possibility of kaempferols regulating plasmodesmata opening via callose deposition. Alternatively, upregulated transcription of callose synthases could be an attempt to control an excessive opening of plasmodesmata. In Arabidopsis seeds, the vascular bundle terminates at the end of the funiculus and releases its content into the chalaza, where the cells of the outer integument are connected by plasmodesmata, representing a symplastic extension of the funicular phloem (Stadler et al, 2005). Therefore, enhanced opening in *tt7* would mean enhanced sugar flux though the outer integument, which could activate starch synthesis. A complementary explanation for the *tt7* starch-excess phenotype could be the inhibition of starch-degrading cytosolic α-glucosidases such as DPE2, given that ability of kaempferol to inhibit these enzymes (Peng et al, 2016). Under this hypothesis, starch degradation in the seed coat would be enzymatically blocked, explaining the higher expression in *tt7* starch-degrading genes as an ineffective attempt by the cellular machinery to react to the high levels of starch.

Readjustment of growth by alterations of cell differentiation status is a strategy of plants suffering from different stresses. Given that flavonoids are involved in the response to a plethora of stressful agents, they are candidates to impact on this stress-induced redistribution of growth and development (Lazar and Goodman 2006; Jan et al, 2021). Environmental conditions experienced by the mother plant alter the developmental programs leading to the acquisition of longevity and other seed traits (He et al, 2014; Nagel et al, 2015), and most of these conditions alter flavonoid composition (Di Ferdinando et al, 2012). PAs content, for example, changes by environmental signals perceived by the mother plant (Macgregor et al, 2015). Changes in flavonol composition could be a mechanism to modulate enzyme activity which impacts cell trafficking and cell-cell communication. Interestingly, the largest variations in flavonoids found in natural accessions of Arabidopsis are in seed coat-specific flavonols, such as quercetin 3-O-rhamnoside or kaempferol derivatives (Routaboul et al, 2012). The fact that the most promising QTL candidate for controlling kaempferol contents is at the genomic region where the TT7 gene is located (Routaboul et al, 2012), and that TT7 is the only flavonoid biosynthetic gene of the genetic network expressed during acquisition of longevity (Riguetti et al, 2015) suggest that modulation of quercetin to kaempferol ratio could be a signal to regulate this trait via cell wall reinforcements and suberin accumulation, positioning TT7 as an essential player controlling seed coat development and seed longevity.

## ACKNOWLEDGEMENTS

This study was funded by the Spanish Ministry of Science and Education, action BIO2014-52621-R. We thank Dr. Javier Forment from the Bioinformatic Unit at the IBMCP for assistance. We thank Alix Mertens and Juliette de Lhonneux for their help in iodine assays, and Virginia Ariza and Raquel Bertí for contribution in *tt7* phenotyping assays. We also thanks Dr. Dolores Planes and Carmen Ruiz-Pastor for occasional assistance in this study during the Covid-19 pandemic.

## AUTHOR CONTRIBUTION

R.N. and P.A. carried out the experiments; J.G. wrote the manuscript with support from R.N. R.S, E.B. and I.M.; A.H. and I.M. performed lipid polyester measurements. J.G and R.N. conceived the original idea.

## DATA AVAILABILITY

Data have been submitted to GEO repository under the accession number GSE197932

