## Supplemental Information for "Kaempferol over-accumulation in the flavonoid 3’ hydroxylase *tt7* mutant disrupts seed coat outer integument differentiation and compromises seed longevity"

**SUMMARY**

Supplemental Figures 1-13

Supplemental Tables 1-3

Supplemental Methods

**SUPPLEMENTAL FIGURES**

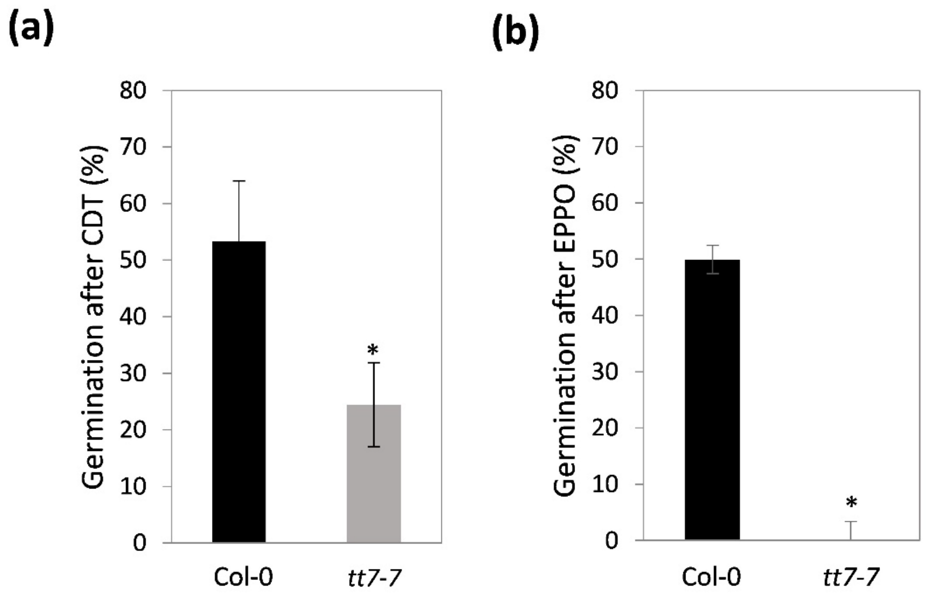


Supplemental Figure 1: Percentage of germination of *tt7-7* and Col-0 seeds after artificial ageing. a) Seeds were incubated during 14 days at 75% RH (CDT) and sown on MS plates. Percentage of germination was recorded after 5 days. b) Seeds from 100%-germinating batches were sown on MS plates after storage for 5 months at 5 bar $O_2$ and 40% RH (Elevated Partial Pressure of Oxygen Assay, EPPO). Percentage of germination was recorded after 5 days. Bars represent the average and standard errors of three replicates with 30 seeds per line. Col-0, Columbia accession. Asterisks indicate significant differences with WT ($P< 0.05$) between samples in two-tailed t-Student test.

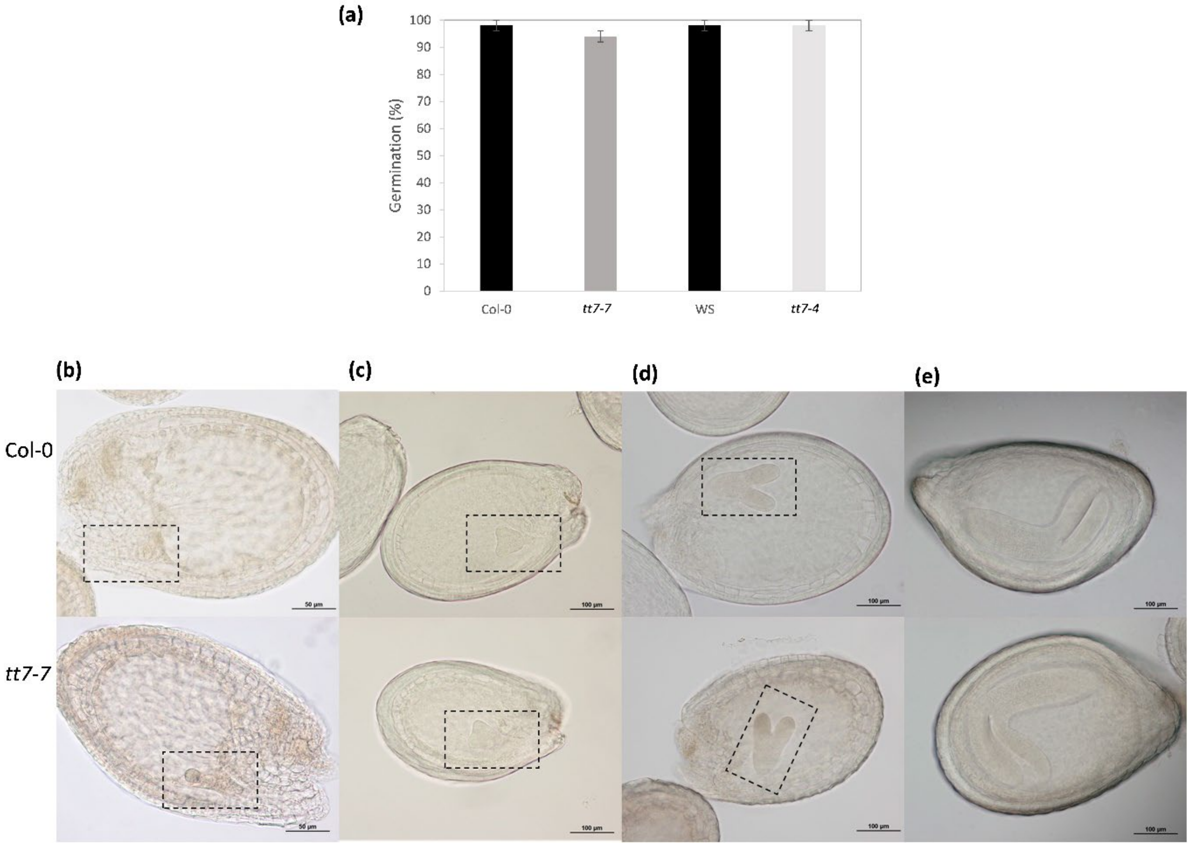


Supplemental Figure 2: <u>*tt7* embryos are viable and develop normally</u>. a) Percentage of germination of Col-0, *tt7-7*, WS and *tt7-4* seeds after growing on MS medium for 7 days. Developing seeds of Col-0 and *tt7-7* mutant were cleared and embryos were observed at 3 (b), 5 (c), 7 (d) and 9 (e) days after pollinization (DAP). Dashed boxes show embryos at globular (d), heart (c) and torpedo stage (d). WS, Wassilewskija. Col-0, Columbia-0.

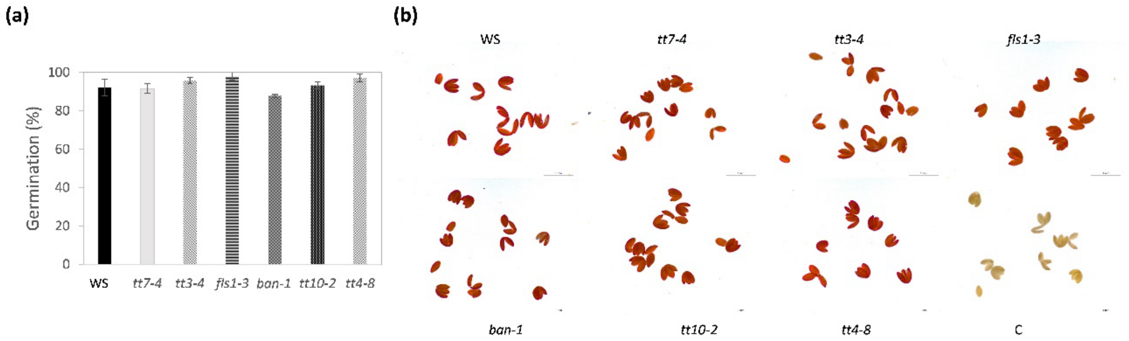


Supplemental Figure 3: *tt* mutant embryos are viable and have similar ability to reduce tetrazolium salts. a) Percentage of germination of different *tt* mutants after 7 days growing in MS medium. Bars represent the mean and standard error of three replicates. b) Embryos from *tt* mutants that were dissected out of the seed coat before incubation in 1% tetrazolium for 20 hours. c) negative control performed with heat-killed seeds (100ºC, 1h). WS, Wassilewskija.

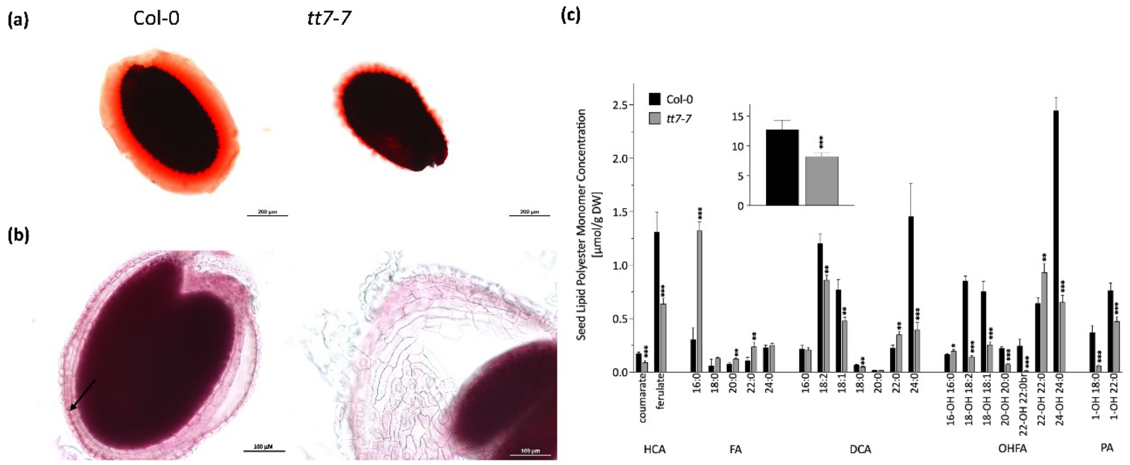


Supplemental Figure 4. <u>*tt7-7* has reduced mucilage and seed coat lipid polyesters</u> a) Representative images of Col-0 and *tt7-7* seeds after staining with ruthenium red during 15 min. Bars: 200 µM. b) Representative images of Col-0 and *tt7-7* seeds stained with Sudan Red 7B. Arrows indicate the palisade layer. Bars, 100 µM. c inset) Total amount of lipid polyester monomers released after depolymerization of Col-0 and *tt7-7* seeds. c) Lipid polyester monomer composition of Col-0 and *tt7-7* mutant seeds. Bars represent the mean content of each identified monomer and error bars represent SD. Asterisks indicate significant differences with WT (*, P< 0.05; **, P< 0.01; ***, P<0.001) between samples in two-tailed t-Student test. br: branched monomer; DCA: dicarboxylic acids; FA: fatty acids; HCA; Hydroxycinnamic acids; OHFA: hydroxyfatty acids; PA: primary fatty alcohols; Col-0: Columbia 0.

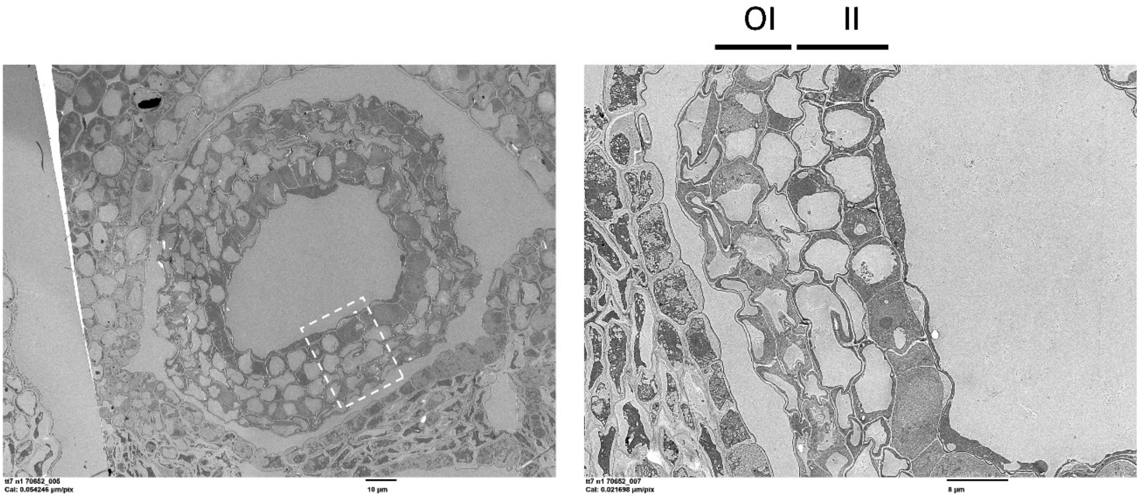


Supplemental Figure 5: <u>*tt7* integuments are well-developed at 3 DAP</u>. TEM images of a representative seed coat of *tt7-4* mutant. White box indicates the area amplified in the right panel. Left panel scale bar, 10 µm; Right panel scale bar, 8 µm; OI: Outer integument; II: Inner integument.

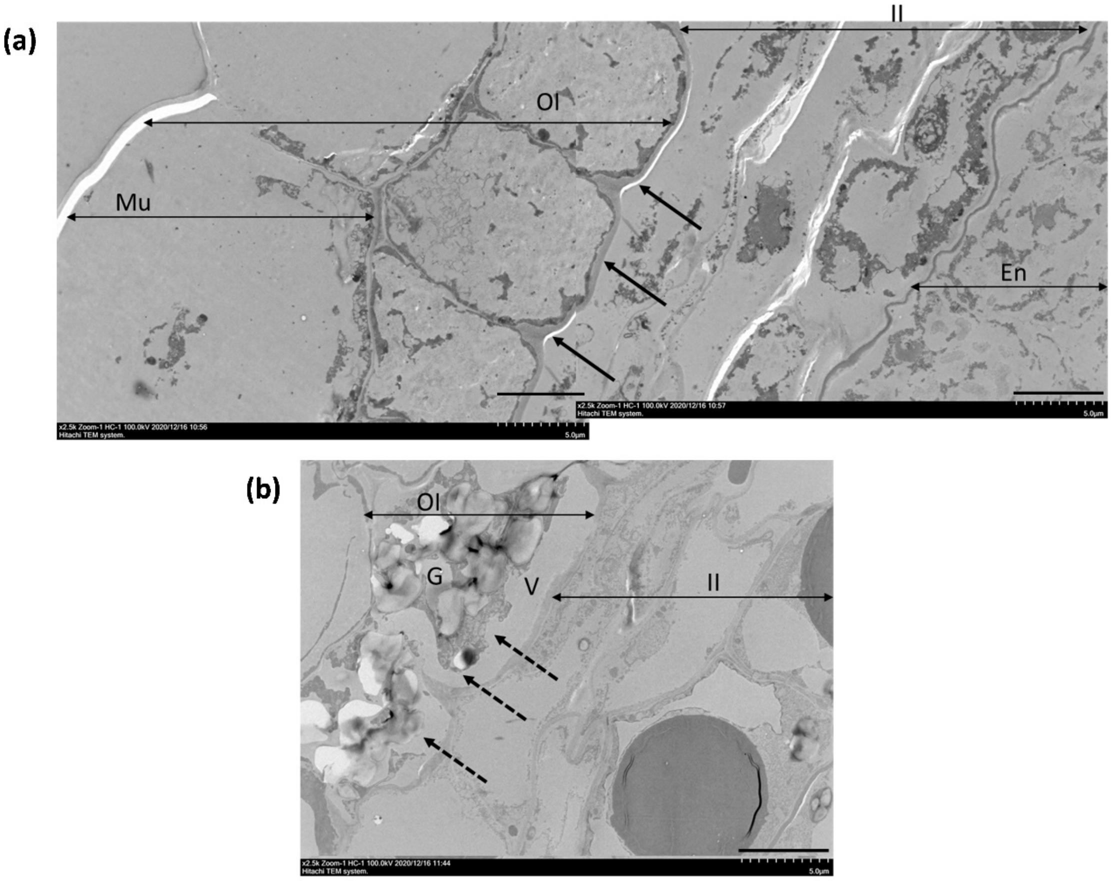


Supplemental Figure 6. Detailed TEM images of WS (a) and *tt7-4* (b) seed coat at 9 DAP. Arrows show the presence of a reinforced cell wall in wild-type seed coat in the inner side of the sub-epidermal layer. Dashed arrows show the absence of reinforced cell wall in *tt7-4* seed coats. Scale bars: 5 µm; II: Inner integument, OI: Outer integument, Mu: Mucilage, G: granules, V:vacuole; WS: Wassilevskija.

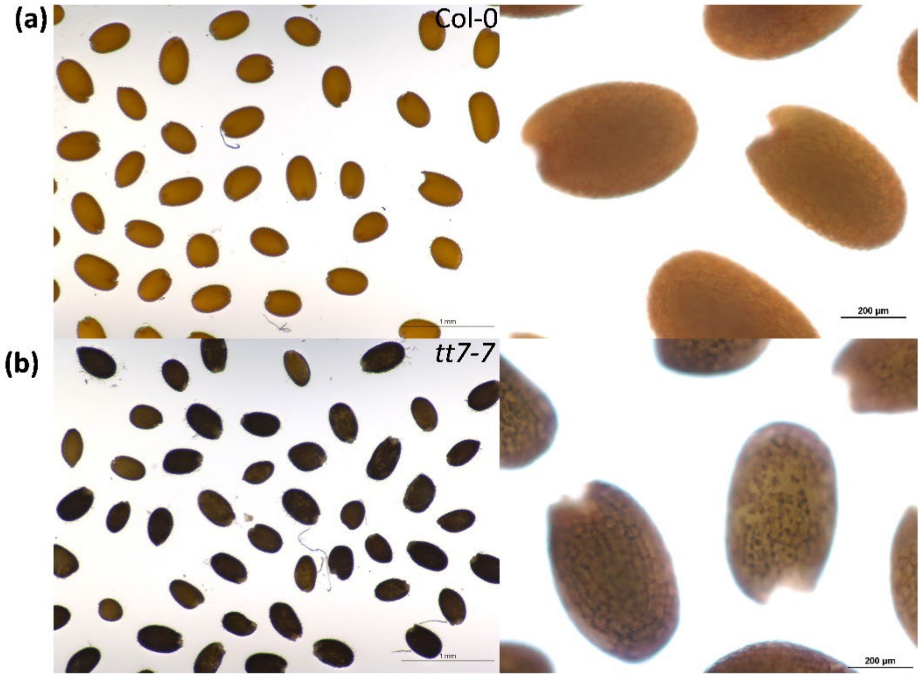


Supplemental figure 7: <u>*tt7-7* mutant seeds accumulate starch.</u> a) Image of Col-0 and *tt7-7* dry seeds after iodine staining. Scale bar, 1mm. b) Amplified view of Col-0 and *tt7-7* dry seeds after iodine staining. Scale bar, 200 µm. Col-0: Columbia 0.

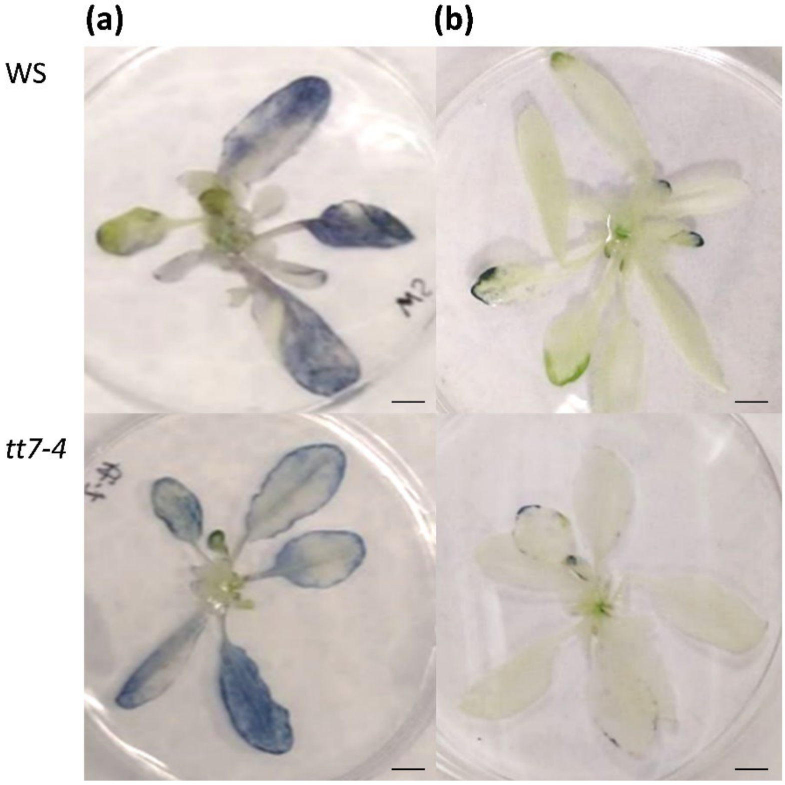


Supplemental Figure 8. <u>*tt7-4* leaves do not accumulate starch</u>. Representative image of rossettes of 18-day-old WS (above) and *tt7-4* plants (below) after iodine staining. a) Leaves after 10h light period; b) Leaves at the end of the dark period. Photoperiod: 8h/16h dark/light. Scale bars: 5 mm; WS: Wassilevskija.

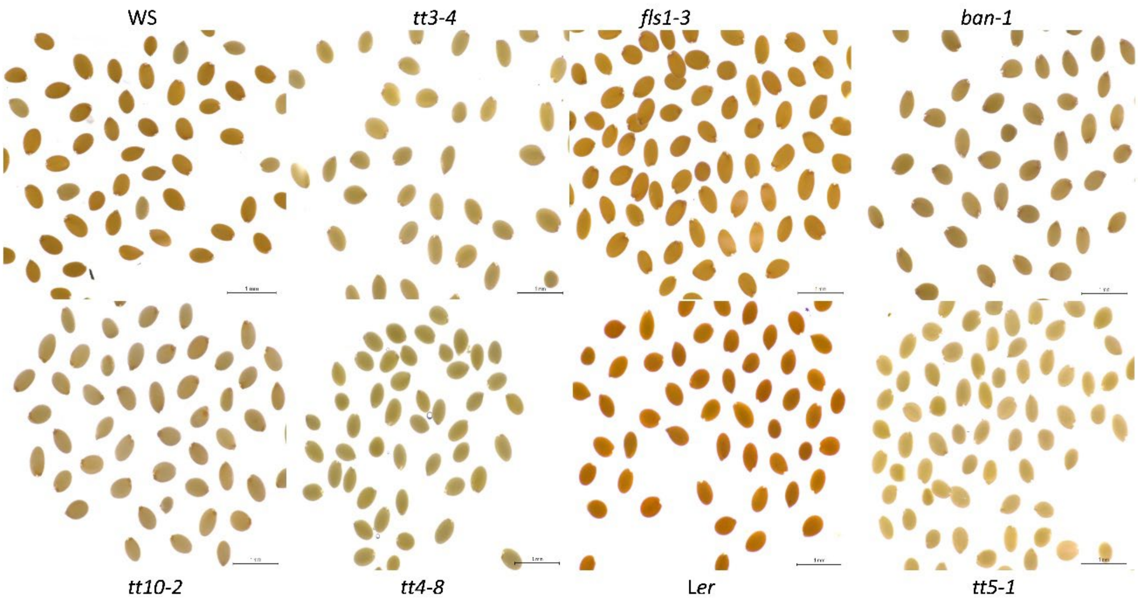


Supplemental Figure 9: *tt7* seed coat starch accumulation is not shared by other *tt* mutants. Representative image of WS and different *tt* mutant dry seeds after iodine staining. Scale bar, 1mm. Ler: Landsberg *erecta*.

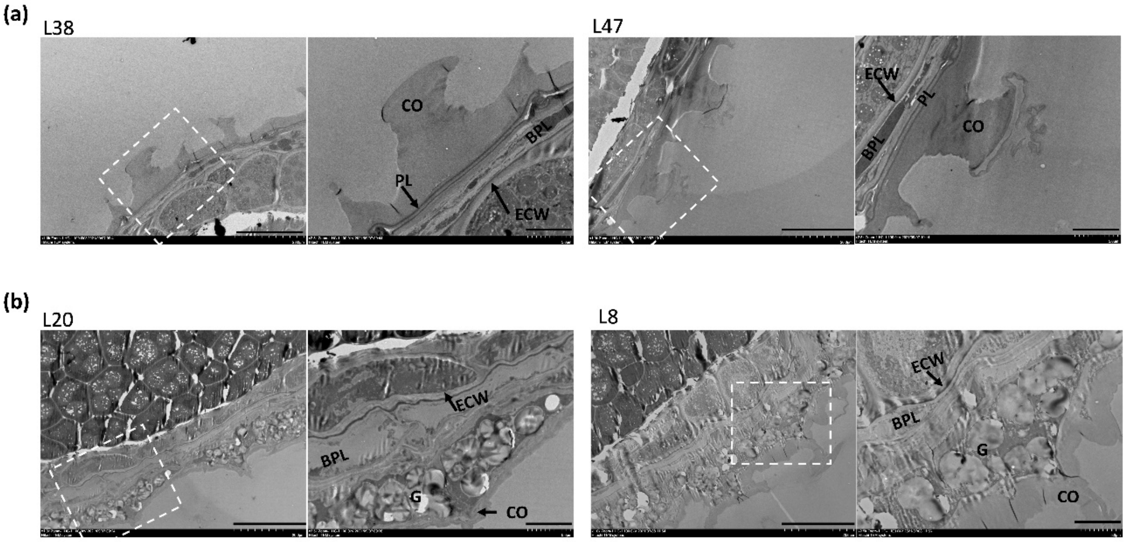


Supplemental Figure 10: Representative TEM images of seed coats of different mutant lines obtained by CRISPR-edition of *TT7* gene in *fls1-3* (L38 and L47) (a), or *tt3-4* (L20 and L8) (b) mutant backgrounds. White boxes indicate the area amplified in right panels. Scale bars in left panels: 20 µm; Scale bars in right panels, 5 µm; ECW: Endosperm cell wall; BPL: Brown pigment layer; PL: Palisade layer; CO: columnella, G: Starch granules

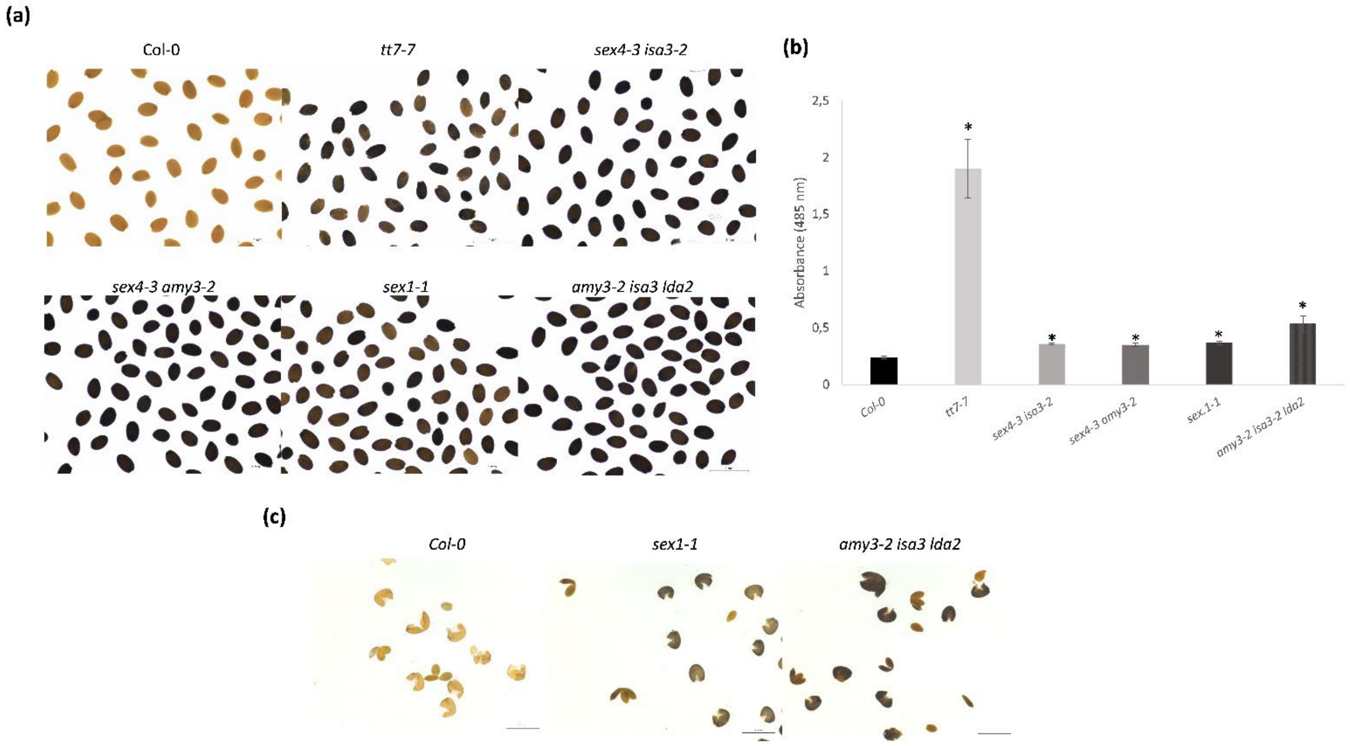


Supplemental Figure 11: Starch accumulation and seed permeability of starch-accumulating mutants. a) Representative image of dry seeds of starch-accumulating mutants after iodine staining. b) Wild-type (Col-0) and mutant seeds were incubated during 2 days in 1% tetrazolium at 30ºC and then formazan was extracted and quantified. c) Iodine staining of separate mature seed coat and embryos of starch-accumulating mutants. Asterisks indicate significant differences with WT (P< 0.05) between samples in two-tailed t-Student test. Col-0: Columbia 0. Scale bar, 1mm

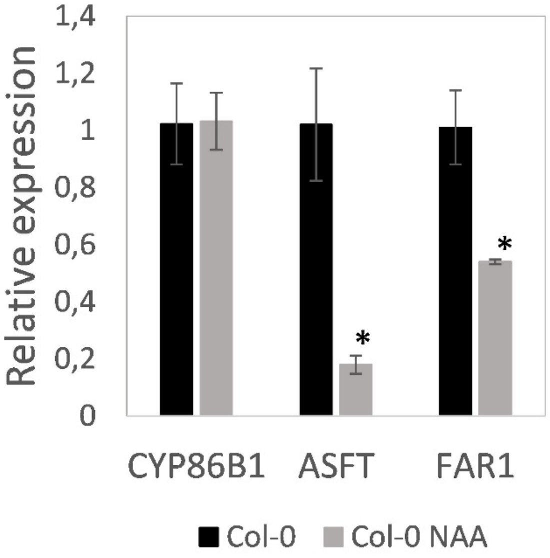


Supplemental Figure 12. Expression of suberin biosynthesis genes after NAA treatment. Expression levels of suberin biosynthesis genes was determined by real-time quantitative PCR in developing seeds (6-10 DAP) of Col-0 plants or Col-0 plants treated with NAA (Col-0 NAA). Developing siliques were sprayed with 50 µM NAA and material was taken 4 h after the treatment. Data are the mean of three replicates. Expression for every gene in Col-0 is set to 1 in the y-axis and relative expression in Col-0 NAA is shown. Significantly differing from wild type at P<0.05 (*) using student´s t-test. Col-0=Columbia 0.

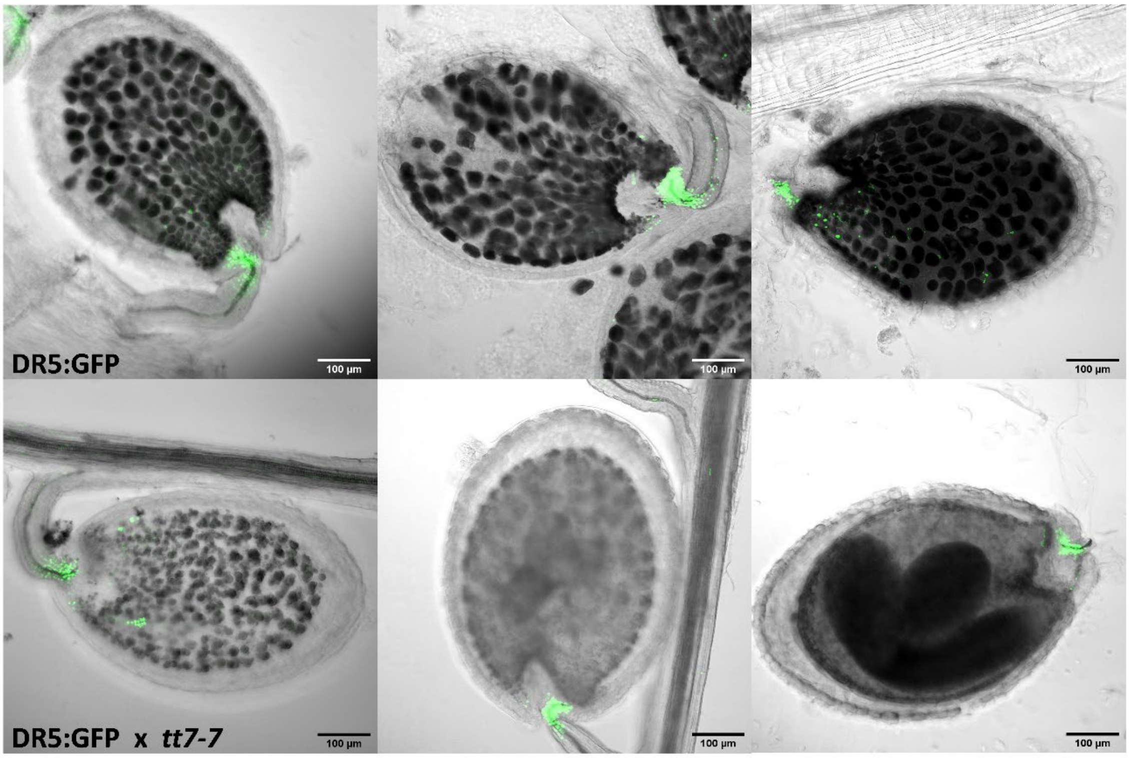


Supplemental Figure 13: Representative confocal images of DR5:GFP (upper panel) and DR5:GFR x *tt7-7* (bottom panel) seeds, 7 (a), 9 (b) and 11 (c ) days after development. Scale bar 100 µm.

**SUPPLEMENTAL TABLES**

Supplemental Table 1: Differentially-expressed genes between 8 DAP wild-type and *tt7-7* seeds.

Supplemental Table 2: Gene ontology categories enriched on differentially-expressed genes between wild-type and *tt7* 8 DAP seeds.

Supplemental Table 3: Primers used in this study

### SUPPLEMENTAL METHODS

#### Lipid Polyesters Analysis

Seed samples (approximately 0.1 g per replicate, 4 replicates per genotype) were incubated in hot isopropanol at 85°C for 15 minutes and ground with a Polytron. The tissue was further delipidated by shaking in different solvents, namely isopropanol, CH3OH:ClCH3 [1:2] and CH3OH:ClCH3 [2:1]) (Jenkin and Molina, 2015); after each incubation samples were centrifuged and the solvent discarded. The cell wall-enriched residues were dried under vacuum for a week, and subsequently depolymerized by NaOMe-catalyzed transmethylation, with the addition of 1 mg g-1 DW each of pentadecanol methyl heptadecanoate (17:0 ME) and pentadecalactone as internal standards (Molina et al., 2006). The depolymerization products were extracted with CH2Cl2 and derivatized by treatment with with pyridine/acetic anhydride. Derivatized samples were evaporated to dryness, resuspended in 500 µL hexane and analyzed by GC-MS on a TRACE 1300 Thermo Scientific gas chromatograph with a ISQ Single Quadrupole mass spectrometer detector. Split injection (5:1 ratio, 310°C) was used with TG-5MS column (Thermo Scientific; 30-m length, 0.25-mm i.d., and 0.25-mm film thickness). The oven temperature was programmed from 140 °C to 310 °C at 3 °C/min and increased to a 310 °C at a rate of 10 °C/min, with a final 10 min-hold at 310 °C. The helium flow rate was set at 1.5 mL/min. The mass spectrometer was run in scan mode over 40-600 amu (electron impact ionization). Monomers were quantified by comparison of each peak area to that of an internal standard.

#### *tt7* editing experiments

Two sgRNAs (5'-N20NGG-3') for *TT7* editing were designed to generate loss-of-function mutations using CRISPR-P 2.0 (http://crispr.hzau.edu.cn/cgi-bin/CRISPR2/CRISPR). The two sgRNAs were designed on the first exon to make a small fragment deletion (153 bp), as shown in Figure 1. After cloning into the pHEE401 binary vector, sequence integrity was confirmed by Sanger sequencing using the sequencing primers detailed in Supplemental Table 3, and the positive vectors harbouring the expression cassettes of the two sgRNAs and zCas9, was introduced into Agrobacterium.

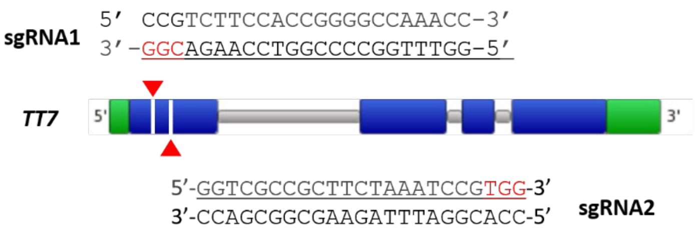


Figure 1. sgRNA sequences designed for loss-of-function mutation in *TT7.* The sgRNAs are designed in the first exon to make a small fragment delection (153b). Blue boxes represent exons, grey lines introns and green boxes 5' and 3'UTRs. Red letters indicate protospacer adjacent motif (PAM) sequences. Red triangles show Cas9 cleavage sites.

**Identification of TT7 knockout transgenic plants**

The transgenic T0 seeds were sown on the MS medium containing 25 mg/L hygromycin at 4 °C for three days in dark and transferred to a growing chamber at 22°C under 16-h-light/8-h-dark photoperiod. After two weeks, Hyg-resistant seedlings (T1) were potted in soil until they grown to 4–6 leaf period. Genomic DNA was extracted from T1 transgenic plant leaves using CTAB method and a PCR with primers surrounding the target sites of TT7 (shown in Figure 2) was performed to identify homozygous tt7 mutants. The expected PCR product size from WT plants is 590 bp, but for tt7 mutants, with a 153 bp deletion, PCR would amplify a fragment of 437 bp (Figure 2). T2 seeds obtained from homozygous T1 mutants were used in our assays.

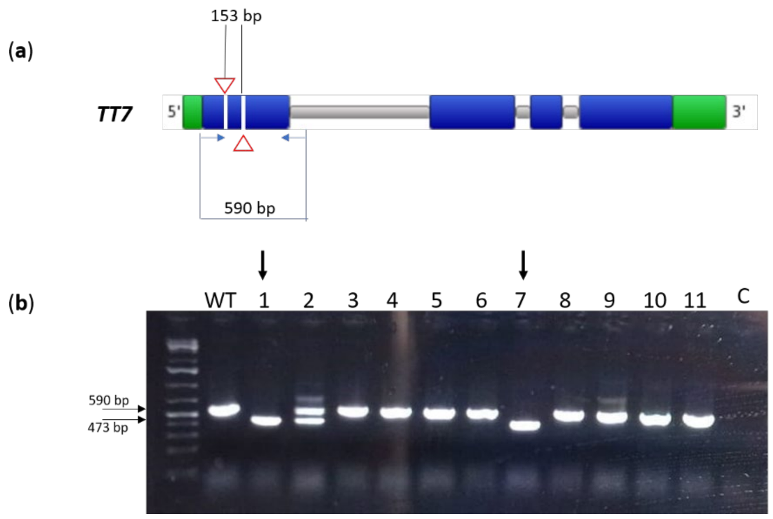


Figure 2. Selection of homozygous *tt7* CRISPR-edited plants by PCR. (a) Scheme of the TT7 gene showing localization of the primers used for PCR (blue arrows), that amplify a fragment of 590bp in a WT and 473 bp in a *tt7* mutant, which harbors a 153 bp deletion.

Red triangles show sites for DNA cleveage. (b) Image of a representative electrophoresis gel after amplifying the TT7 locus of eleven candidates (plants transformed with EC1.2::Cas9 and two guide RNAs directed to TT7). A band of 590 bp is expected in the wild type (first well), whereas a band of 473 bp is expected in the mutant. Arrows indicate two homozygous mutant lines. C: negative control. WT: wild-type plant.

**RNA seq analysis**

Twenty-million paired-ends 50pb reads per library (40 million in total) were sequenced. After adaptor removal and low-quality trimming of raw reads with cutadapt (Martin, 2011), clean reads were quality assessed with FastQC (Andrews, 2010) and mapped to the TAIR10 Arabidopsis thaliana genome using HISAT2 (Kim et al, 2015). Gene counts were then obtained with htseq-count (Anders et al, 2015) and used for differential expression analysis with DESeq2 (Love et al, 2014). Significative genes (padjusted <0,05 and expression in tt7 at least two-fold higher or lower than in the wild type) were used for functional analysis using AgriGOv2 (Tian et al, 2017). Redundant GO terms were removed and remaining GO terms were visualised using ReviGO (Supek et al, 2011).

**Transmission electron microscopy**

Samples were hydrated and chemically fixed with Karnovsky's solution. Seeds were then washed in phosphate buffer and post-fixed in 2% osmium tetroxide. Samples were dehydrated through a graded ethanol series and embedded in LR resin for 24h in 55°C. All samples were sectioned (60-90 nm) using an Ultracut Leica UC6 with a diamond knives (Diatome). Sections for each sample were examined and imaged with a JEM-1010 transmission electron microscope.

**Clearing and visualizing Arabidopsis embryos**

3-9 DAP siliques were fixed with 2 ml 100% EtOH: acetic acid (9:1) solution and vacuum was applied for 15 m (twice). Samples were incubated for 2h at room temperature, with fresh fixation solution. Then, samples were maintained for 30 m in 96% EtOH and 30 m in 70% EtOH. Chloral hydrate was added and maintained for 3-5 days in the dark. For embryo visualization, seeds were separated from siliques in a slide with glycerol/water (2:1) and observed in a Nikon Eclipse E600 microscope.

Andrews, S. (2010). FastQC: A Quality Control Tool for High Throughput Sequence Data [Online].Available at:http://www.bioinformatics.babraham.ac.uk/projects/fastqc/

Anders S, Pyl PT, Huber W. HTSeq--a Python framework to work with high-throughput sequencing data. Bioinformatics. 2015 Jan 15;31(2):166-9. doi: 10.1093/bioinformatics/btu638.

Jenkin, S., Molina, I., 2015. Isolation and compositional analysis of plant cuticle lipid polyester monomers. J. Vis. Exp. 1–10. https://doi.org/10.3791/53386

Kim D, Langmead B and Salzberg SL. HISAT: a fast-spliced aligner with low memory requirements. Nat Methods 12, 357–360 (2015). https://doi.org/10.1038/nmeth.3317

Love MI, Huber W, Anders S (2014). "Moderated estimation of fold change and dispersion for RNA-seq data with DESeq2." Genome Biology, 15, 550. doi: 10.1186/s13059-014-0550-8.

Martin, M. Cutadapt removes adapter sequences from high-throughput sequencing reads. EMBnet.journal, [S.l.], v. 17, n. 1, p. pp. 10-12, may 2011. ISSN 2226-6089.

Molina, I., Bonaventure, G., Ohlrogge, J., Pollard, M., 2006. The lipid polyester composition of Arabidopsis thaliana and Brassica napus seeds. Phytochemistry 67, 2597–2610. https://doi.org/10.1016/j.phytochem.2006.09.011

Supek F, Bošnjak M, Škunca N, Šmuc T. REVIGO summarizes and visualizes long lists of gene ontology terms. PLoS One. 2011;6(7):e21800. doi: 10.1371/journal.pone.0021800

Tian T, Liu Y, Yan H, You Q, Yi X, Du Z, Xu W, Su Z. agriGO v2.0: a GO analysis toolkit for the agricultural community, 2017 update. Nucleic Acids Res. 2017 Jul 3;45(W1):W122-W129. doi: 10.1093/nar/gkx382
